# Selective Control of Parasitic Nematodes Using Bioactivated Nematicides

**DOI:** 10.1101/2022.03.11.483960

**Authors:** Andrew R. Burns, Rachel J. Ross, Megan Kitner, Jonathan R. Volpatti, Aditya S. Vaidya, Emily Puumala, Bruna M. Palmeira, Elizabeth M. Redman, Jamie Snider, Sagar Marwah, Sai W. Chung, Margaret H. MacDonald, Jens Tiefenbach, Chun Hu, Qi Xiao, Constance A. M. Finney, Henry M. Krause, Sonya A. MacParland, Igor Stagljar, John S. Gilleard, Leah E. Cowen, Susan L. F. Meyer, Sean R. Cutler, James J. Dowling, Mark Lautens, Inga Zasada, Peter J. Roy

## Abstract

Parasitic nematodes are a major threat to global food security, particularly as the world amasses 10 billion people amidst limited arable land. Most traditional nematicides have been banned due to poor nematode-selectivity, leaving farmers with inadequate controls. Here, we use the model nematode *Caenorhabditis elegans* to identify a family of selective imidazothiazole nematicides, called selectivins, that undergo cytochrome p450-dependent bioactivation exclusively in nematodes. At low parts-per-million concentrations, selectivins perform comparably well with commercial nematicides to control root infection by *Meloidogyne incognita* – the world’s most destructive plant-parasitic nematode. Tests against a wide range of phylogenetically diverse non-target systems demonstrate that selectivins are more nematode-selective than nearly all marketed nematicides. Thus, selectivins are first-in-class bioactivated nematode controls that provide efficacy as well as much-needed nematode selectivity.

## MAIN TEXT

Global food demand will be increasingly difficult to meet as the human population approaches 10 billion people^1–4^. There is a scarcity of arable land for agricultural expansion, and land conversion efforts are constrained by social and ecological factors. Maximizing production from currently cultivated land will be crucial to ensure global food security^1–4^.

Farmers rely on agrochemicals to maximize yields by controlling crop pathogens^4, 5^. Plant-parasitic nematodes (PPNs) are especially destructive pathogens that cause over $100 billion in crop losses every year^6, 7^. Synthetic nematicides have played an essential role in PPN control for decades, however concerns over environmental toxicity and human safety have justifiably prompted bans on the most commonly used nematicides^8, 9^. The gutting of available nematode controls is stark – of the 20 key nematicides used in the 20^th^ century, only 4 are currently approved for use in the European Union, and only 2 are used in the United States without restriction^9–11^. Though warranted, these withdrawals leave farmers with limited options, and no control measures for several PPNs^8, 12^.

Despite the need for more selective PPN controls, only six non-fumigant synthetic nematicides have been developed in the past twenty-five years^9, 13^. One of these “next-generation” nematicides (iprodione) has already been banned in Europe, due in part to its carcinogenic potential and risk to aquatic life^10^, and the market release of another (tioxazafen) has been postponed due to skin irritation reported by handlers. There is a pressing need for new nematicides with improved selectivity.

### Selective imidazothiazole nematicides

From a library of uncharacterized compounds that disrupt the growth of the free-living nematode *Caenorhabditis elegans*^14^ we identified three novel imidazothiazole-containing molecules that selectively kill nematodes (Fig. 1). At low micromolar concentrations, these <u>selectiv</u>e imidazothiazole <u>n</u>ematicides, which we call selectivin-A, -B, and -C, kill all four of the free-living nematode species that we culture in the lab, as well as the eggs of a nematode parasite of cattle, and the infective juveniles of two distinct species of root-knot nematode that infect crops. These seven nematode species span five genera and two evolutionary clades suggesting that diverse nematode species, including parasites, are susceptible to the selectivins. By contrast, these compounds exhibit no activity against fungi, plants, insects, fish, and human cells at concentrations that readily kill nematodes. Furthermore, mice given a daily oral dose of 50 mg/kg selectivin-A for five days showed no obvious pathologies relative to untreated controls. Broad-spectrum nematicidal activity has never been described for any compound containing the selectivin 6-phenylimidazo[2,1-*b*]thiazole core scaffold, and we are the first to demonstrate efficacy against plant-parasitic nematodes for this structural class. Thus, the selectivins represent an entirely new chemistry for the development of selective PPN controls.

**Fig. 1.**
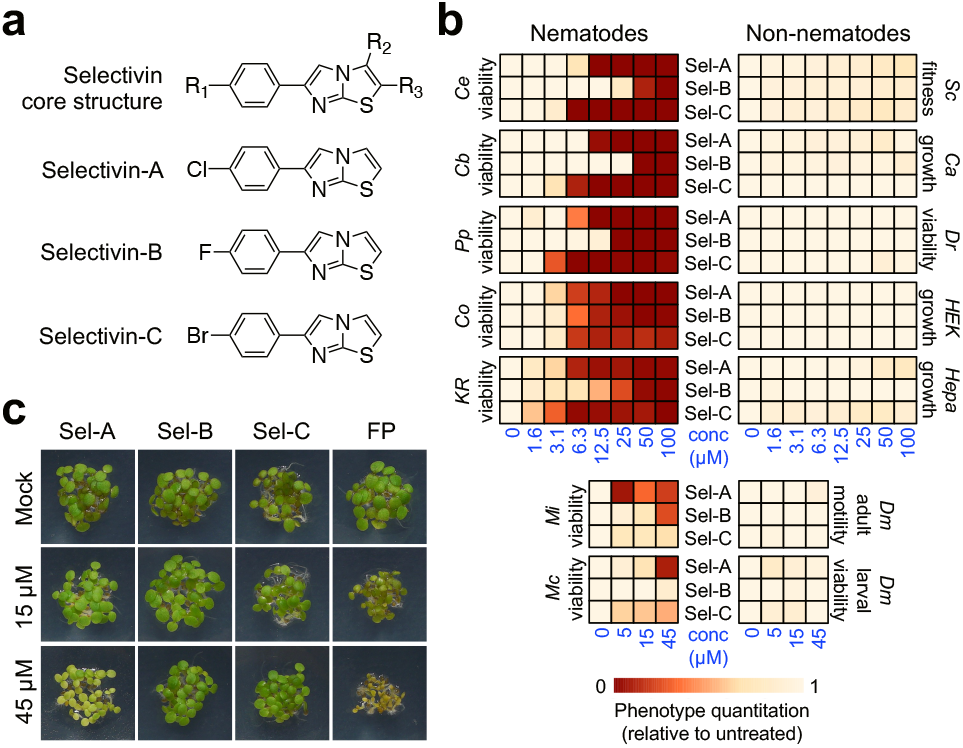
Selective imidazothiazole nematicides (selectivins). **a**, The selectivin core scaffold, and structures of selectivin-A, -B, and -C. **b,** Selectivin (Sel) dose-response for seven nematode species and six non-nematode model systems. The colour-coded scale denotes the phenotypic outcome of selectivin treatment, relative to untreated control. *Ce*, *Caenorhabditis elegans* (clade V); *Caenorhabditis briggsae* (clade V); *Pp*, *Pristionchus pacificus* (clade V); *KR*, *Rhabditophanes sp.* KR3021 (clade IV); *Mi*, *Meloidogyne incognita* (clade IV); *Mc*, *Meloidogyne chitwoodi* (clade IV); *Sc*, *Saccharomyces cerevisiae* (fungus); *Ca*, *Candida albicans* (fungus); *Dr*, *Danio rerio* (fish); *HEK*, HEK 293 cells (human); *Hepa*, HepaRG cells (human); *Dm*, *Drosophila melanogaster* (insect). **c,** *Arabidopsis thaliana* (plant) seedling establishment and greening in response to selectivins and the commercial nematicide fluopyram (FP).

### Selectivins have a new mode-of-action

As a first step towards understanding the selectivin mode-of-action, we tested selectivin-A and selectivin-C against eight *C. elegans* mutant strains that are each specifically resistant to one of the major anthelmintic or nematicide classes (Extended Data Table 1)^14–22^. We found that none of the mutants are selectivin-resistant, suggesting that selectivins have a distinct mode-of-action compared with commercial agents. Selectivins are structurally similar to the commercial anthelmintic levamisole, which is used to treat livestock infected with parasitic nematodes^23^. The selectivins and levamisole both contain imidazothiazole rings as part of their core structure^23^; however, the selectivins differ from levamisole in that their imidazothiazole ring is unsaturated, whereas levamisole contains a partially saturated tetrahydroimidazothiazole ring. Despite this structural similarity, levamisole-resistant mutants are fully sensitive to the selectivins, indicating that these compounds have a unique mode-of-action.

To further establish the distinctiveness of the selectivins, we performed a forward genetic screen for *C. elegans* mutants (generated from random mutagenesis) that resist selectivin-induced lethality. This approach has previously yielded large numbers of mutants that specifically resist many of the major classes of anthelmintics and nematicides that are used commercially^14, 15, 17, 19, 20, 22, 24^. Despite screening over 10 million mutagenized genomes, selectivin-resistant worms could not be generated (Extended Data Table 2). This result reinforces the conclusion that selectivins kill nematodes via a mechanism that is distinct from traditional nematode controls, and suggests that parasitic nematodes may not develop resistance in the field.

### Selectivins are bioactivated nematicides

Previous work with *C. elegans* has shown that metabolites of nematode-lethal compounds often accumulate in worm tissue^25^. It is therefore possible that a nematicide could be transformed from an innocuous ‘parent’ molecule into a metabolic product that is lethal to worms. Microsomal cytochrome p450 (CYP) enzymes are commonly involved in the oxidative biotransformation of inert pro-drugs into bioactive metabolites^26^. The *C. elegans* genome encodes 76 microsomal CYP enzymes (Supplementary Table 1)^27^, many of which are known to metabolize drugs^28^. To test whether selectivins are bioactivated by *C. elegans*, we capitalized on previous work that achieved broad disruption of microsomal CYPs by compromising EMB-8 activity during larval development, bypassing its essential role during embryogenesis^28^. EMB-8 is the *C. elegans* p450 oxidoreductase (POR) enzyme that is a necessary cofactor of all microsomal CYPs^28, 29^. Consistent with the selectivins being bioactivated nematicides, we found that EMB-8/POR knockdown rendered *C. elegans* completely resistant to selectivin-induced lethality (Fig. 2a). By contrast, the compound wact-11, which we have previously shown does not require bioactivation for activity^14^, kills worms in an EMB-8/POR-independent manner.

**Fig. 2.**
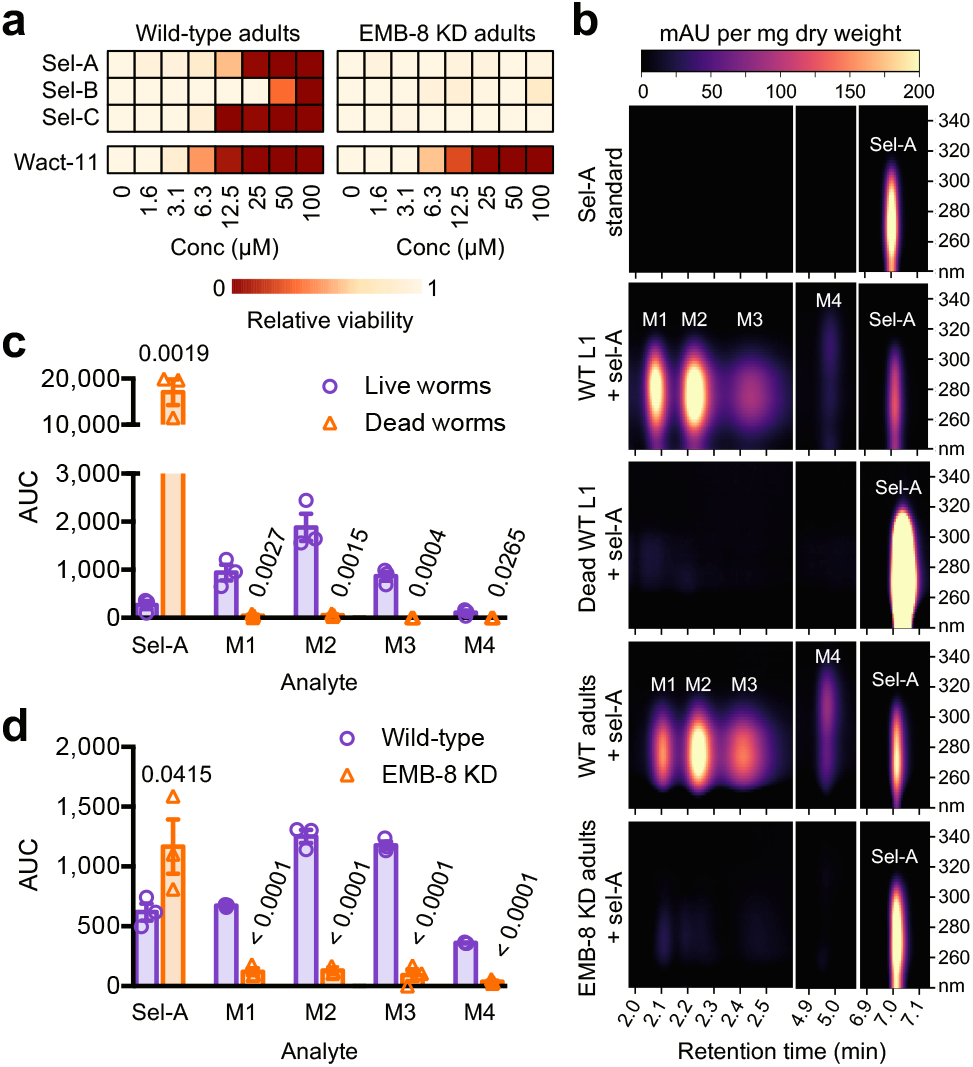
Selectivins are bioactivated nematicides. **a,** Selectivin and wact-11 dose-response for wild-type (WT) and EMB-8 knockdown adult worms. **b,** HPLC chromatograms of lysates from live and dead L1-stage worms, as well as WT and EMB-8 knockdown adults, treated with 100 µM selectivin-A (sel-A). The chromatograms shown are the product of dry weight normalization of raw chromatograms, followed by subtraction of absorbance intensity from paired untreated controls. Absorbance intensity is in milli-absorbance units (mAU) per milligram of dry weight, and absorbance wavelength (y-axis) is in nanometers (nm). The top chromatogram is the sel-A standard injected directly onto the column, and is not biomass-normalized or background-corrected. Unmodified sel-A and metabolites M1-M4 are indicated. **c,** Quantification of unmodified sel-A and M1-M4 in the lysates of live and dead L1s. **d,** Quantification of sel-A and M1-M4 in the lysates of WT and EMB-8 KD adults. In both **c** and **d** AUC is the area under the curve for the given analyte peak, calculated at 260 nm for M1-M3 and at 300 nm for M4. Error bars are SEM. For each analyte, P-values were obtained from unpaired Student’s t-tests comparing the means of live and dead worms (**c**), or WT and EMB-8 KD worms (**d**).

To identify selectivin metabolites, we incubated both young adult and L1 stage worms in the presence or absence of 100 µM selectivin-A for 6 hours and then processed whole worm lysates by reversed-phase HPLC coupled with an absorbance detector (Fig. 2b). We identified five obvious peaks in the selectivin-A-treated lysates that are absent from the untreated controls. The peak that elutes at 7 minutes invariably has the same retention time and absorbance spectrum as the pure selectivin-A standard injected directly onto the column. This peak is likely unmodified selectivin-A parent molecule that persists in worm tissue following the incubation. The remaining four peaks elute earlier in the HPLC run relative to selectivin-A, suggesting that they are relatively less lipophilic. These peaks are absent from the selectivin-A standard, and they are not found in the lysates of dead worms incubated in selectivin-A, consistent with them being *bona fide* selectivin-A metabolites and not simply accumulated contaminants or spontaneous oxidation products (Fig. 2b, c). We have named these presumptive selectivin-A metabolites M1, M2, M3, and M4.

Consistent with our hypothesis that selectivin-A is bioactivated, the abundance of all four metabolites in worm tissue is dramatically reduced upon EMB-8/POR knockdown, whereas the abundance of the unmodified selectivin-A parent increases by almost 2-fold (Fig. 2b, d). Unmodified selectivin-A is unlikely to be the active agent *in vivo* because EMB-8/POR knockdown worms survive treatment with 100 µM selectivin-A despite the internal concentration being nearly twice as high as that found in wild-type worms (Fig. 2a, d). These results indicate that selectivins are bioactivated pro-nematicides.

### Bioactivated to a toxic electrophile

To identify the toxic selectivin metabolite(s), we collected HPLC fractions containing the four selectivin-A metabolites and identified their structures by electrospray ionization mass spectrometry (ESI-MS) followed by MS-MS fragmentation (Fig. 3a-d). Abundant masses in the M1, M2, and M3 fractions, and respective MS-MS fragmentations, are consistent with singly protonated ions of γ-glutamylcysteine, glutathione, and cysteine conjugates of a selectivin-A sulfoxide metabolite, respectively (Fig. 3a-c; Extended Data Table 3). Alkylating lysis did not result in a reduction in the abundance of the conjugates in worm lysates, suggesting that they form *in vivo* (Extended Data Fig. 1). These data reveal a canonical detoxification pathway in which glutathione is conjugated to the sulfoxide metabolite, thereby sequestering it from cellular nucleophiles, and is then converted to γ-glutamylcysteine and cysteine, resulting in metabolites that are readily excretable (Fig. 3e)^30–33^. Notably, accumulation of the low-molecular-weight thiol adducts in worm tissue suggests that sulfoxide metabolite production outpaces the worms’ capacity for detoxification.

**Fig. 3.**
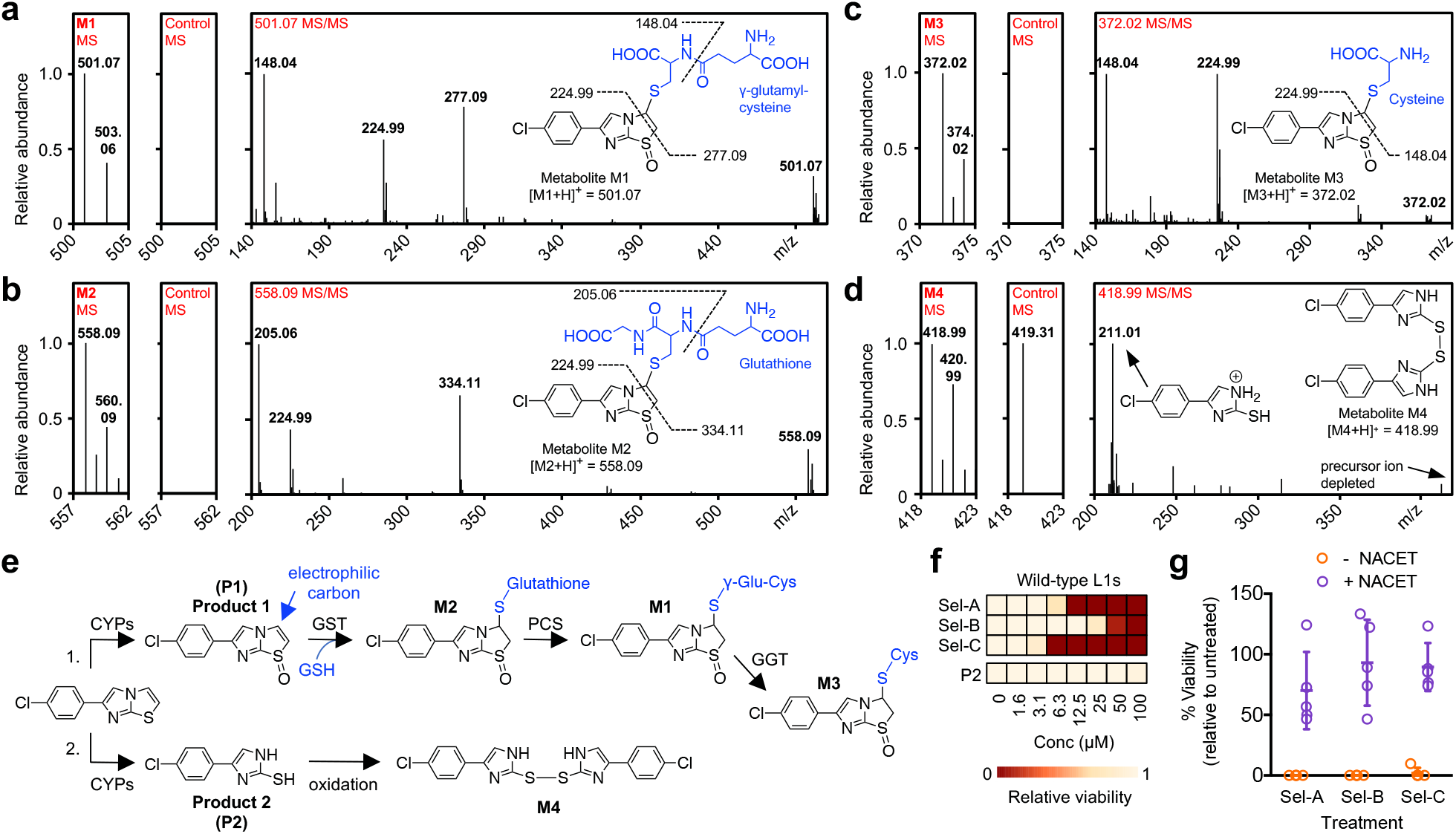
Selectivins are bioactivated to a toxic electrophile. **a-d,** Mass spectrometry (MS) data for metabolite M1-M4 HPLC fractions and untreated control fractions, as well as MS/MS fragmentation data for selected masses. Structures of γ-glutamylcysteine (γ-Glu-Cys) (**a**), glutathione (**c**), and cysteine (**c**) conjugates of a selectivin-A electrophilic sulfoxide metabolite are shown, along with fragmentations that produce masses found in the MS/MS plots. The proposed M4 disulfide structure is shown in (**d**), along with the structure of the protonated imidazole-thiol monomer that results from disulfide bond cleavage. **e,** Schematic of the two major selectivin-A metabolism pathways: 1. S-oxidation produces Product 1 (P1), which conjugates with glutathione (GSH) either spontaneously or via glutathione-S-transferase (GST), and is further processed to γ-Glu-Cys and cysteine conjugates via phytochelatin synthase (PCS) and γ-glutamyl transferase (GGT), respectively; 2. Imidazothiazole ring opening to produce Product 2 (P2), which oxidizes to form the disulfide M4. Cytochrome p450 enzymes (CYPs) likely mediate ring opening and S-oxidation. **f,** Dose-response for *C. elegans* L1s treated with selectivins and P2. **g,** Viability of L1 worms treated with selectivins, alone or in combination with 2.5 mM NACET. Error bars are SD.

Our analysis of the M4 fraction revealed a 419.00 mass that is consistent with a disulfide formed between two monomers of 4-(4-chlorophenyl)-1*H*-imidazole-2-thiol, and MS-MS fragmentation produced a mass consistent with the monomer (Fig. 3d; Extended Data Table 3). Alkylating lysis results in alkylated monomer, but not the disulfide, indicating that the imidazole-thiol monomer is likely a metabolite of selectivin-A that forms *in vivo*, whereas the disulfide forms post-lysis (Extended Data Fig. 1; Extended Data Table 3). Thus, CYP-mediated metabolism of selectivin-A results in two metabolic products. Product 1 is an electrophilic sulfoxide and product 2 is an imidazole-thiol metabolite (Fig. 3e).

Either product 1 or product 2 could kill nematodes. Dose-response analysis with product 2, which is commercially available, revealed that it is unable to kill worms up to a concentration of 100 µM, despite readily accumulating in worm tissue (Fig. 3f, Extended Data Fig. 2). Hence, product 2 is not responsible for selectivin lethality. We could not directly test whether product 1 induces lethality because it could not be synthesized, despite trying numerous oxidation schemes (see Methods). Instead, we tested whether the exogenous addition of N-acetylcysteine ethyl ester (NACET), which is enzymatically converted into cysteine for use in the synthesis of new glutathione^34^, can suppress the lethality induced by the selectivins. NACET has been used to block the damage caused by acetaminophen overdose, which results from the buildup of a reactive electrophilic metabolite that depletes glutathione and reacts with proteins causing hepatotoxicity^34^. We reasoned that product 1 may kill nematodes by a similar mechanism. Indeed, 2.5 mM NACET suppresses the lethality induced by the selectivins, consistent with product 1 being the active nematicidal agent *in vivo* (Fig. 3g). These data indicate that selectivin metabolism produces a toxic electrophile that kills nematodes.

### Bioactivation is nematode specific

Given their selective nematicidal activity, we hypothesized that bioactivation of selectivins may be phylogenetically restricted to nematodes. Indeed, microsomal CYP families are not well-conserved between nematodes and other phyla (Supplementary Table 1)^7, 27, 35–41^. For example, none of the microsomal CYP families that overlap between *C. elegans* and the plant-parasitic nematode *Meloidogyne incognita* are found in vertebrates, insects, plants, or fungi. By contrast, five CYP families are conserved within diverse nematode species, and CYPs from at least two of these families metabolize drugs^28^.

To test the phylogenetic specificity of selectivin bioactivation, we analysed selectivin-A metabolites in both the lysates and incubation buffers of four distinct nematode species spanning three genera and two evolutionary clades, as well as in three non-nematode species from diverse phyla (Extended Data Fig. 3a, b). The non-nematode species tested were the yeast *Saccharomyces cerevisiae*, the fly *Drosophila melanogaster*, and the fish *Danio rerio*. We found that the low-molecular-weight thiol conjugates of the sulfoxide metabolite are abundantly produced by all four nematode species (Extended Data Fig. 3a-c, Extended Data Tables 3 and 4). By contrast, the three non-nematode species produce significantly less of the conjugates relative to nematodes (Extended Data Fig. 3a-c). Indeed, the conjugates are undetectable by mass spectrometry in yeast lysate, and only the cysteine conjugate can be detected at very low levels in the lysates of flies (Extended Data Table 4). Furthermore, the glutathione conjugate (M2) readily accumulates in the tissues of all four nematode species over the course of the incubation, but not in the tissues of the non-nematode species, suggesting that abundant sulfoxide metabolite production and oversaturation of the detoxification pathway is common to nematodes but not to distinct phyla (Extended Data Fig. 3a, d). Notably, we found that tissue accumulation of the selectivin-A parent compound in the non-nematode species was comparable to, or greater than, accumulation in *C. elegans* tissue (Extended Data Fig. 3a, e). This result suggests that poor bioavailability is unlikely to account for the lower levels of metabolite production by non-nematode species, and provides further evidence that the selectivins require bioactivation to kill. Taken together, these data indicate that distinct nematode species from disparate evolutionary clades are capable of robustly bioactivating the selectivins, and that bioactivation is phylogenetically restricted to nematodes.

### A lead nematicide to control PPNs

Encouraged by the selectivity, broad-spectrum action, and novel mechanism of the selectivins, we performed an analog series to identify a lead nematicide capable of effectively controlling root infection by the root-knot nematode *Meloidogyne incognita* – arguably the world’s most destructive crop pathogen^42^. Eleven analogs were included in the series, along with all four of the marketed next-generation soil-applied synthetic nematicides for which powder is commercially available: fluensulfone, fluopyram, iprodione, and tioxazafen. Selectivin analogs were either purchasable or synthesized using the route shown in Extended Data Fig. 4 (see Methods). Tests were performed against *M. incognita* in its soil environment with a tomato host plant (Fig. 4a). Soil-based assays are critical for establishing the real-world utility of candidate nematicides since many compounds that are effective *in vitro* lose activity in the soil due to poor mobility and/or degradation^43^. We assayed compounds by adding 45 µM of aqueous compound and between 1,000 and 2,500 infective J2 nematode larvae to 90 grams of potted soil. The final soil concentration is approximately 2.5 parts-per-million (ppm) or 4.5 kg per hectare (kg/ha), which is comparable to the application rates of commercial nematicides^43, 44^. 24 hours later, we planted 2 to 3-week old tomato seedlings in the soil. Eight weeks later, the effectiveness of the compounds at reducing eggs in the roots was calculated relative to an untreated control. Fluensulfone and fluopyram demonstrated nearly 100% effectiveness and were the best performing nematicides. Consistent with previous reports, iprodione was the least effective commercial nematicide^13^. Selectivin-E was the most effective analog tested (56% effective) and performed comparably well with tioxazafen (57% effective) and outperformed iprodione (38% effective). Of note, we challenged the plants with a nematode density that is at least 2-fold greater than what is considered high density in fields^45, 46^, which may account for why some of the nematicides tested had incomplete efficacies in this assay. Treatment with the vast majority of selectivins, including selectivin-E, increases root weights relative to untreated control (Extended Data Fig. 5), likely by reducing nematode burden. Encouragingly, none of the selectivins tested were obviously toxic to tomato plants, suggesting that this family of compounds is not generally phytotoxic.

**Fig. 4.**
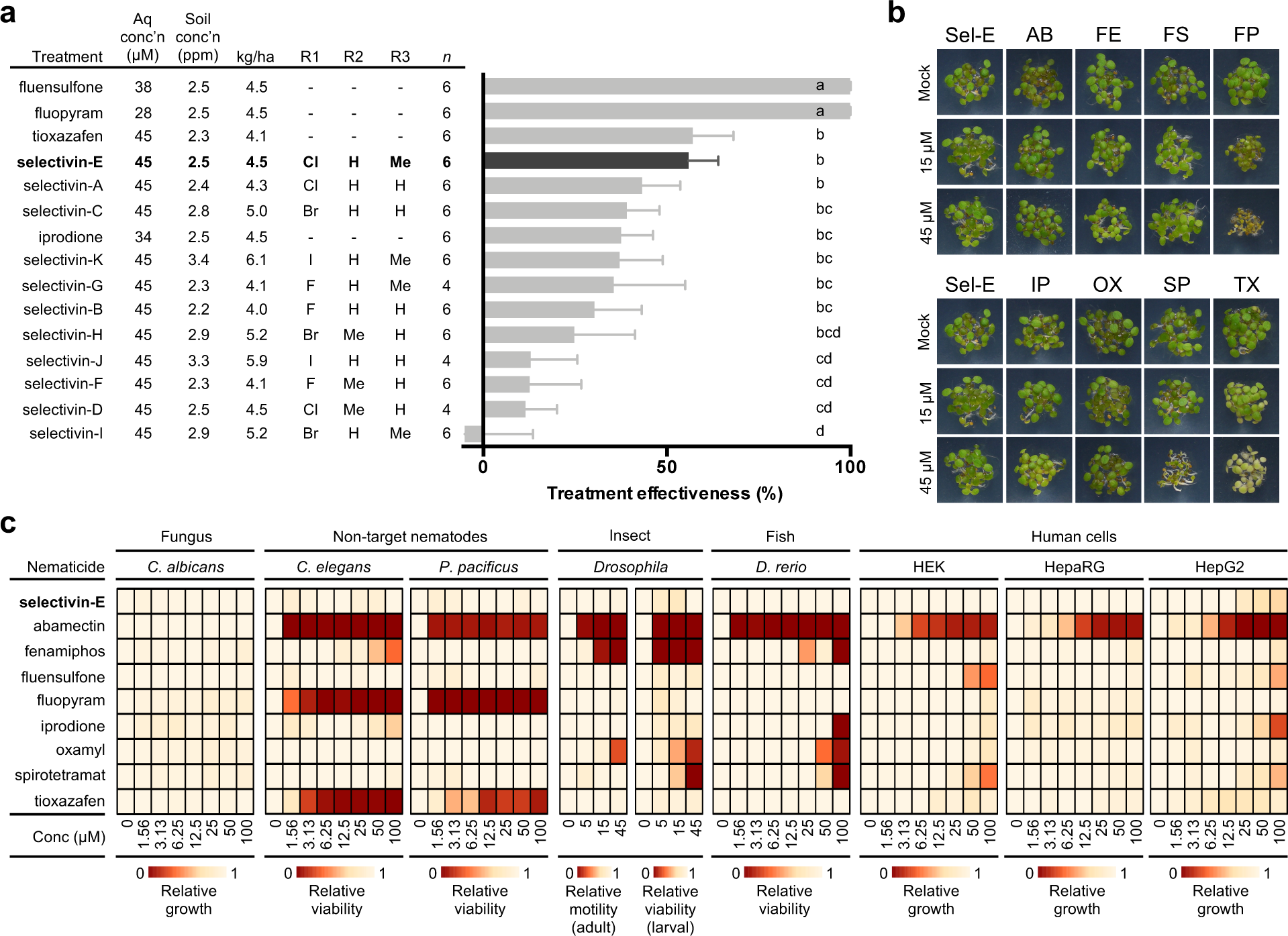
Selectivin-E is a lead nematicide for plant-parasitic nematode control. **a,** Percent effectiveness of 11 selectivin analogs and 4 commercial nematicides at preventing tomato plant root infection by *M. incognita* (see Methods). For each analog, aqueous molar concentration, parts-per-million soil concentration, and kilograms-per-hectare values are indicated. The R-groups for each analog are also indicated (see Fig. 1a for the selectivin core structure and R-group positions). Means followed by an uncorrected Fisher’s LSD analysis (P > 0.05 is not significant). **b,** *Arabidopsis thaliana* seedling establishment and greening in response to selectivin-E and eight commercial nematicides. AB, abamectin; FE, fenamiphos; FS, fluensulfone; FP, fluopyram; IP, iprodione; OX, oxamyl; SP, spirotetramat; TX, tioxazafen. **c,** Dose-response of phylogenetically diverse non-target systems treated with selectivin-E and eight commercial nematicides. The phenotypic outcome of each treatment, relative to untreated control, is denoted by the colour-coded scales.

To assess the selectivity of selectivin-E for PPNs we tested it against a panel of phylogenetically diverse non-target systems including plants, fungi, free-living nematodes, insects, fish, and three different human cell types (Fig. 4b, c). Two of the human cell types are HepaRG and HepG2 cells, which are human liver progenitor cells that are well-established models for assessing drug metabolism and bioactivation in human liver – inactivity in both cell types suggests a low potential for drug-induced liver injury^47^. We also included all of the newly developed synthetic nematicides in our tests, as well as the natural product abamectin, and the organophosphate/carbamate nematicides fenamiphos and oxamyl, which is still in use. Selectivin-E has the best selectivity profile compared with all of the nematicides tested. Selectivin-E is inactive against all of the non-target systems up to 100 µM, which was the highest concentration assayed. Notably, selectivin-E is non-lethal to the two free-living nematodes tested – albeit developmental delay is observed for *C. elegans* at 100 µM – suggesting that selectivin-E may be relatively innocuous to soil-beneficial free-living nematodes. By contrast, every commercial nematicide tested had unwanted toxicity in at least one of the non-target assays.

Furthermore, selectivin-E is lipophilic (cLogP = 4.038) and only moderately water soluble (LogSw = -4.88), suggesting that it will not readily leach into ground water and is unlikely to be taken up systemically by plants^48^, thereby limiting residues in fruits and other plant products ingested by consumers. Taken together, these data suggest that our best performing lead nematicide, selectivin-E, can effectively and selectively control nematode pests.

## Discussion

Here, we report our discovery of the nematicidal utility and mode-of-action of the selectivins – the first nematicide class known to kill via metabolic bioactivation in nematodes. From a small set of analogs, we have identified a promising lead compound that can effectively control the notorious plant-parasitic nematode (PPN) *Meloidogyne incognita*. Given the broad-spectrum activity of the selectivins against diverse nematode species, and their innocuity against non-nematode phyla (including plants), this novel class of nematicides has the potential to selectively control a wide-variety of PPNs. Unlike several new nematicides on the market, which are repurposed fungicides or insecticides^9, 13^, the selectivins are “true nematicides” that target nematodes selectively.

Crop protection nematicides are deployed in metric tonne quantities, therefore they must have scalable synthetic routes and be cheap to produce. The synthetic scheme we chose to produce selectivin analogs satisfies many of the criteria for a large-scale industrial process. Selectivin synthesis by this method requires only two steps, two easily accessible solvents, moderate reaction temperatures and pressures, and no transition metal catalysts. Furthermore, the synthesis of selectivin-E makes use of inexpensive, easily accessible, and relatively non-toxic starting reagents. The yields we report are two-step yields – intermediates were not purified or isolated between steps, which is advantageous for large-scale synthesis.

Our work demonstrates for the first time that worm-bioactivated compounds can be broadly effective against disparate worm species. This result is perhaps unexpected since broad-spectrum activity of a worm-activated compound will depend upon low interspecies and intraspecies variability in enzyme expression and activity. Indeed, drug metabolism commonly varies between individuals of the same species^49^. Thus, it would not be surprising if a pro-nematicide approach was overlooked by industrial discovery programmes. Although we focus on PPN control, this approach could also be applied to anthelmintics used to treat animal parasites. We highlight that the free-living nematode *C. elegans*, which has largely been abandoned by industry as a discovery platform^50^, was the system that revealed the selectivins. Our work thus reinforces the idea that *C. elegans* can be a useful model system to identify and characterize promising nematicidal leads.

## METHODS

### Free-living nematode strains and culture methods

The *Caenorhabditis elegans* wild-type N2 strain and the MJ69 strain harbouring the temperature-sensitive *emb-8(hc69)* allele were obtained from the *C. elegans* Genetics Center (CGC, University of Minnesota). *Caenorhabditis briggsae* strain AF16 and *Pristionchus pacificus* strain PS312 were both obtained from the CGC. *Rhabditophanes sp. KR3021* was obtained from Marie-Anne Félix (Institute of Biology of the Ecole Normale Supérieure (IBENS), Paris, France). The following anthelmintic/nematicide-resistant *C. elegans* strains were used for experiments: VC731 (*unc-63(ok1075)*), PR1152 (*cha-1(p1152)*), RP2674 *(sdhc-1(tr393)*), CB3474 (*ben-1(e1880)*), RB2119 (*acr-23(ok2804)*), NM1968 (*slo-1(js379)*), CF1038 (*daf-16(mu86)*), DA1316 (*avr-14(ad1302);avr-15(vu227);glc-1(pk54)*). With the exception of RP2674, which was obtained from a genetic screen performed in our lab, all of these strains were obtained from the CGC. With the exception of *C. elegans* strain MJ69, all nematode species and strains were cultured and propagated at 20°C using standard techniques^51^. *C. elegans* strain MJ69 was propagated at the permissible temperature of 15°C. We found that using solid NGM agar plates with thin bacterial lawns and high nematode densities promoted growth of *Rhabditophanes sp.* KR3021.

### Commercial chemical sources

Selectivin-A, -B, -G, -J, -K, and tioxazafen were purchased from ChemBridge Corporation. Selectivin-C was purchased from Vitas-M and MilliporeSigma. Selectivin-D was purchased from MolPort. 4-(4-chlorophenyl)-1*H*-imidazole-2-thiol was purchased from Life Chemicals Incorporated. Abamectin, fenamiphos, fluopyram, iprodione, oxamyl, spirotetramat, and iodoacetamide were purchased from MilliporeSigma. Fluensulfone was purchased from Cayman Chemical.

### C. elegans dose-response experiments

An HB101 bacterial suspension in liquid NGM was prepared by concentrating a saturated overnight HB101 culture 2-fold with liquid NGM (see ref. 14 for NGM recipe). Forty microliters of this bacterial suspension was added to each well of a 96-well flat-bottom culture plate, after which approximately 25 synchronized L1 worms, in 10 µl of M9 buffer (see ref. 52 for the recipe), were added to each well. The synchronized L1 worms were obtained from an embryo preparation performed the previous day (see ref. 52 for the protocol). For the L1 assays, 0.5 µl of chemical solution (or DMSO alone) was immediately added to the wells using a multichannel pipette; the final DMSO concentration was 1% (v/v). The culture plate was sealed with parafilm and placed in a box with several wet paper towels to prevent evaporation. The worms were incubated for 3 days at 20°C with shaking at 200 rpm and the number of viable animals was counted. A dead worm was considered any worm that failed to move after vigorous agitation of the plate, and that appeared morphologically “dead”, i.e. clear appearance and unresolved internal structures. Although the counts were performed after 3 days of incubation in the chemical, it was noted that the L1s were dead within 24 hours of the addition of the chemicals. For the wild-type young adult dose-response assays, HT115 *E. coli* carrying the empty dsRNA expressing vector L4440 was used in place of HB101 and the worms were cultured at 25°C as opposed to 20°C for the entirety of the experiment. The HT115 suspension was made by concentrating a bacterial culture, with an OD_600_ of 0.8, five-fold with liquid NGM containing 1 mM IPTG and 100 µg/ml carbenicillin. The HT115 cells were induced with 1 mM IPTG for one hour at 20°C before concentrating with NGM. Synchronized L1 worms were grown for 2 days until they reached young adulthood, at which point chemical was added to the wells by multichannel pipette. Viability was scored one day later. The EMB-8 knockdown dose-response assays were performed identically to the wild-type young adult dose-response experiments, except the MJ69 strain was used in place of the wild-type strain, and worms were fed HT115(DE3) *E. coli* harbouring an RNAi feeding vector expressing dsRNA targeting the *emb-8* gene.

The dose-response experiments for the anthelmintic/nematicide-resistant mutants were carried out as described for the L1 dose-response assays. One notable exception is the aldicarb-resistant strain PR1152. This strain grows slowly, and so the viability counts were performed 5 days after addition of the chemical, as opposed to 3 days, to allow the DMSO control worms to reach adulthood. At least three biological replicates were performed for each dose-response assay. For each biological replicate, two technical replicates were performed and the numbers of viable animals for each technical replicate were combined (i.e. ∼ 50 worms assayed per concentration). The number of viable worms at each concentration was divided by the corresponding DMSO control value to give the “relative viability” for each concentration. The “relative viability” values were then averaged across the biological replicates. Where appropriate, LC_50_ values were calculated using Graphpad Prism – the concentration values were log-transformed and a four-parameter logistic curve was fitted to the dose-response data by non-linear regression, from which the LC_50_ values were extracted.

### C. briggsae, P. pacificus, and Rhab. sp. KR3021 dose-response experiments

Dose-response assays and quantification for *C. briggsae*, *P. pacificus*, and *Rhabditophanes sp.* KR3021, were carried out as described for the *C. elegans* L1 dose-response assays; however, the selectivin-induced phenotypes in *P. pacificus* and *Rhabditophanes*, even at the highest concentrations, were a combination of lethality and larval arrest with severe sickness. Therefore, for these dose-response assays, the number of animals that reached the L2 stage or older was quantified, as opposed to the number of viable worms. The arrested animals appeared very sick, and would likely die before reaching reproductive adulthood. Therefore, this arrested phenotype was considered to be practically analogous to death.

### NACET experiments

Saturated HB101 bacterial cultures were concentrated 2-fold with either regular liquid NGM or liquid NGM containing 2.5 mM NACET. In the wells of a 96-well culture plate, 25 synchronized L1 worms in 10 µL of M9 buffer were added to forty microliters of the bacterial suspensions. The synchronized L1 worms were obtained from an embryo preparation performed the previous day (see ref. 52 for the protocol). The culture plate was sealed with parafilm and placed in a box with several wet paper towels. The worms were incubated for 4 hours at 20°C, after which 100 µM of nematicide, or DMSO alone, was added independently to NACET-containing or NACET-free wells. The plates were re-sealed with parafilm, replaced in the box with wet paper towels, and allowed to incubate for an additional 24 hours at 20°C before assessing lethality. Lethality was assessed as described above for the *C. elegans* L1 dose-response assays. 2.5 mM NACET was chosen because it is the highest concentration of NACET that can be used without dramatically slowing worm growth or killing the worms outright. Two technical replicates were performed for each condition, and the number of living worms was summed across the two replicates. The number of viable worms for each condition was divided by the corresponding DMSO control value to give the “relative viability” for each condition. This was repeated five separate times to give five independent biological replicates.

### Cooperia oncophora dose-response experiments

Fresh cattle faeces containing eggs of an ivermectin-resistant strain of *C. oncophora* were kindly supplied by Dr. Doug Colwell and Dawn Gray (Lethbridge Research Station, Agriculture and Agri-Food Canada). Established methods were used to carry out the experimental cattle infections^53^, and these methods were approved by the Lethbridge AAFC Animal Care committee and conducted under animal use license ACC1407. Cattle faeces containing *C. oncophora* eggs were stored anaerobically at room temperature for a maximum of 6 days before use. Eggs were isolated from faeces using a standard saturated salt flotation method^54^ immediately before the egg hatch assay. 80 µl of distilled and deionized water was added to each well of a 96-well culture plate, and then 1 µl of chemical at the appropriate concentration in DMSO was added to each well using a multichannel pipette.

Approximately 50 eggs were added per well in 20 µl of water for a final volume of 100 µl in each well; the final DMSO concentration was 1% (v/v). The eggs were incubated in the chemicals for 2 days at room temperature, after which hatching was stopped by the addition of 1 µl iodine tincture to each well. The number of hatched larvae was counted at each concentration, and eggs that failed to hatch were scored as dead. “Relative viability” values were calculated by dividing the fraction of eggs that hatched at each concentration by the fraction of eggs that hatched in the corresponding DMSO control well. Two biological replicates were performed for each dose-response experiment, and the relative viability values were averaged across the biological replicates. The average hatch rate for the DMSO control wells was greater than 93% for both biological replicates.

### Meloidogyne spp. in vitro dose-response experiments

*Meloidogyne incognita* (Kofoid & White) Chitwood Race 1 (originally isolated in Maryland) maintained on pepper (*Capsicum annuum* L.) cv. PA-136 and *M. chitwoodi* (originally isolated from Washington) maintained on tomato (*Solanum lycopersicum*) were grown in a greenhouse as previously described^55, 56^. Infective second-stage juveniles (J2) were collected as described in ref. 57. The microwell dose-response experiments were conducted in 96-well polystyrene plates, similarly to previously described protocols^55, 58^. In brief, each well received approximately 35-50 J2s in 10 µL of sterile distilled water (SDW), followed by addition of 190 µL SDW, or of 190 µL of SDW containing dissolved chemical or DMSO alone. The chemicals were tested at 5, 15, and 45 µM concentrations, and the final concentration of DMSO in each well was 0.5% (v/v). The wells were covered with a plastic adhesive strip, and the lids of the plates were sealed with parafilm. The nematodes were incubated in chemicals for 2 days at 26°C, at which point the J2 were rinsed twice with SDW, and incubated in the second SDW rinse for an additional day. Post-incubation, the fraction of viable nematodes at each concentration was calculated by dividing the number of motile nematodes by the total number of nematodes in the well. “Relative viability” was calculated by dividing the fraction of viable nematodes at a given treatment concentration by the fraction of viable nematodes in the untreated DMSO control sample. The experiment was conducted twice, with three wells per treatment in each trial. Average “relative viability” was calculated across six replicates.

### Saccharomyces cerevisiae (budding yeast) dose-response experiments

A saturated culture of the yeast strain RY0568 was diluted to an OD_600_ of 0.015 with fresh YPD media (see ref. 59 for recipe). 100 µL of this dilute yeast suspension was added to each well of a 96-well plate. The yeast were grown for 4 hours at 30°C with shaking at 140 rpm. Using a multichannel pipette, 1 µL of chemical solution was added to each well to achieve the desired final concentrations. The final DMSO concentration was 1% (v/v). The microwell plate was then loaded into a TECAN plate reader set at 30°C. The OD_600_ of each well was measured over an 18-hour period, and the plate was shaken intermittently throughout the run. The areas under the resultant growth curves were calculated using R scripts adapted from those found in the MESS package. The area under the curve at each concentration of a dose-response assay was divided by the area under the curve for the corresponding DMSO control, resulting in a “relative fitness” value for each concentration tested. Three biological replicates were performed for each dose-response experiment, and the relative fitness values were averaged across the three replicates.

### Candida albicans dose-response experiments

Compound potency against *C. albicans* (SN95) was assessed using two-fold dose-response assays following a standard protocol. Briefly, YPD medium was inoculated with ∼1 x 10^3^ cells/mL from saturated overnight cultures. Assays were performed in 384-well, flat-bottom microtiter plates (Corning) in a final volume of 40 µL/well. Each drug was added to wells in a two-fold concentration gradient from 100 µM to 0 µM. Plates were then incubated in the dark at 30 °C under static conditions for 48 hours. Endpoint growth was quantified by measuring optical density (OD) at 600 nm using a spectrophotometer (Molecular Devices) and the results were corrected for background media. Relative fungal growth for each compound treatment was defined by normalizing to the levels observed in untreated controls. All dose-response assays were performed in technical triplicate and relative growth levels were averaged across replicates.

### Danio rerio (zebrafish) dose-response experiments

Fish were maintained at 28.5°C on a 14/10 hour light/dark cycle and staged according to hours post fertilization (hpf). For each biological replicate, eggs from AB wild-type fish were collected at 4 hpf. At 3 days post fertilization (dpf), embryos were arrayed in 24 well culture plates; 10 per well. 5 µl of chemical dissolved in DMSO at the appropriate concentration was added to 1 mL of E3 media and then vortexed intensively to mix. Water was removed from the embryos in the wells and 1 mL of chemical-treated water was transferred to each of the wells. The DMSO control wells contained DMSO alone. The final DMSO concentration in every well was 0.5% (v/v). The culture plates were sealed with parafilm, wrapped in aluminum foil, and incubated at 28.5°C for 72 hours. After the incubation, viability was assessed. Using a stereomicroscope, lethality was scored either by appearance of whole-body necrosis or the absence of both a heartbeat and touch-evoked response. “Relative viability” was calculated by dividing the number of viable embryos in the treatment wells by the number of viable embryos in the DMSO control well. Three biological replicates were performed for each dose-response experiment, and the relative viability values were averaged across the three replicates.

### HEK cell dose-response experiments

HEK293 cells were seeded into 96-well plates, at 5000 cells per well, in 100 µL total volumes of DMEM/10%FBS/1%PS media and grown overnight at 37°C in the presence of 5% CO_2_. Compounds (0.5 µL volumes from appropriate source plates) were then added to cells, and growth was continued for an additional 48 hours. Following growth, 10 µL of CellTiter-Blue Viability reagent (Promega) was added to each well, and plates were incubated for an additional 4 hours at 37°C in the presence of 5% CO_2_. Fluorescence measurements (560 nm excitation/590 nm emission) were then performed using a CLARIOstar Plate Reader (BMG Labtech) to quantify reagent reduction and estimate cell viability. Fluorescence measurements were corrected for background from medium. “Relative growth” was calculated by dividing corrected fluorescence in the treatment wells by that measured in the corresponding DMSO control well. The final values are an average across at least 3 replicates.

### HepaRG cell dose-response experiments

HepaRG cells (Thermofisher) were seeded in a 96-well plate at 20,000 cells per well in 100 µL of DMEM-F12 (GIBCO), +10% FBS (Invitrogen), 1 x Pen/Strep (Sigma), 2mM L-glutamine (Sigma), 1x ITS-A (GIBCO), 40 ng/ml dexamethasone (BioShop). Subsequently, the media was refreshed, and a dilution series of test compound was added to each well with a final total volume of 100 µL. After 48 hours, Cell Titer Blue viability dye (Promega) was added for 3 hours, and viability was quantified using plate reader as per the manufacturers protocol (Promega). Fluorescence was measured at 560 nm excitation/590 nm emission and corrected for background from the medium. “Relative growth” was calculated by dividing corrected fluorescence in the treatment wells by that measured in the corresponding DMSO control well. The final values are an average across at least 3 replicates.

### HepG2 cell dose-response experiments

HepG2 cells (ATCC, Cat# HB-8065) were counted using a haemocytometer, and diluted to 5 x 10^4^ cells/mL in 100 uL of RPMI-1640 (Sigma) medium supplemented with 10% heat inactivated fetal bovine serum (Gibco) following a standard protocol. Cells were seeded in black, clear-bottom 384-well plates (Corning) to a final density of 2000 cells/well in 40 µL. Cells were incubated at 37°C with 5% CO_2_ for 24 hours. Subsequently, a 2-fold dilution series of test compound was added to seeded cells from 0 µM to 100 µM, and plates were incubated at 37°C with 5% CO_2_ for 72 hours. After 72 hours, Alamar Blue (Invitrogen) was added to the HepG2 cells at a final concentration of 0.5X and plates were incubated at 37°C for 4 hours. Fluorescence was measured at Ex560nm/Em590nm using a TECAN Infinite F200Pro microplate fluorometer and values were corrected for background from the medium. All assays were performed in technical triplicates and in at least two biological replicates. “Relative growth” was calculated by dividing corrected fluorescence in the treatment wells by that measured in the corresponding DMSO control well.

### Drosophila melanogaster (fruit fly) dose-response experiments

Fly food in agar substrate was prepared by mixing together 100 mL of unsulfured molasses, 100 mL of cornmeal, 41.2 g of Baker’s yeast, and 14.8 g of agar into 1400 mL of distilled de-ionized water and boiling for 30 minutes. The media was allowed to cool to 56°C, at which point 5 mL was added by syringe to plastic cylindrical fly vials. 10 µL of chemical, or DMSO alone, was added to the media in each vial. The chemicals were mixed into the media by mechanical mixing using a pipette. The final DMSO concentration was 0.2% (v/v). The media was allowed to solidify at room temperature (∼22°C) overnight. The following day (Day 0), eight pairs of male and female w1118 flies were added to each vial so that there were 16 flies in total per vial. The vials were stored at room temperature for 7 days, at which point the number of motile flies was counted. Fly motility was scored as any observable movement after the vial had been vigorously jostled. “Relative motility” was calculated by dividing the number of motile flies in the treatment vials by the average number of motile flies in two DMSO control vials. On Day 8 the 16 parental flies were removed from the vials and the progeny larvae were allowed to continue to grow and hatch into adult flies. To assess larval viability, hatched flies were counted and discarded on Days 10, 12, 14, 16, 18, and 20. The counts were summed. “Relative viability” was calculated by dividing the number of hatched flies in the treatment vials by the average number of hatched flies in the two DMSO control vials. The final “relative motility” and “relative viability” values are an average across three experimental replicates.

### Arabidopsis thaliana greening experiments

Greening experiments were performed with *Arabidopsis thaliana* seeds of wild type Col-0; seeds were surface sterilized in bleach and plated onto 0.5X MS, 0.5% sucrose agar medium supplemented with compounds of interest at 5, 15 and 45µM concentrations (0.2% DMSO (v/v/)). After 4d of stratification at 4°C, plates were transferred to a growth chamber (16h / 8h, 150 µE/m^2^) and greening recorded after 4 days. Pictures were recorded by camera (SONY a7s) with FE1.8/55 lens (FE 55 mm F1.8 ZA; SEL55F18Z). Experiments were performed in triplicate for each treatment.

### Selectivin-A mouse studies

Female C57BL/6 mice (bred in house, breeding pairs originally purchased from Charles River, Canada) 6-8 weeks of age were used for all experiments. Animal experiments were approved by the University of Calgary’s Animal Care Committee. Infected mice were orally gavaged with 200 third stage *Heligmosomoides polygyrus* larvae (maintained in house. Original stock was a gift from Dr. Allen Shostak, University of Alberta, Canada) and euthanized on day 22 post infection. Each group (treated vs. non-treated) had a minimum of 7 mice. Mice were littermates. Mice were treated orally with 5 daily doses of selectivin-A (50mg/kg resuspended in DMSO). Control mice were given DMSO only as a control. Mice were monitored daily for visible signs of treatment-induced pathology.

### Incubations for HPLC analysis

For the *C. elegans* and *C. briggsae* L1 incubations, synchronized hatchlings were obtained from an embryo preparation of gravid adults^52^. 200,000 hatchlings in 500 µL of M9 buffer (see ref. 52 for the M9 buffer recipe) were treated with either 100 µM selectivin-A, 100 µM 4-(4-chlorophenyl)-1*H*-imidazole-2-thiol (i.e. Product 2), or DMSO alone for control purposes. The final concentration of DMSO in all samples was 1% (v/v). Prior to the incubations, the hatchlings used for the dead worm controls were heat-killed at 95°C for 45 minutes. The incubations were carried out in standard 1.5-mL micro-centrifuge tubes on a nutating shaker, at 20°C for 6 hours. After the 6-hour incubation, the worms were transferred to the wells of a Pall AcroPrep 96-well filter plate (0.45-µm wwPTFE membrane, 1-ml well volume), the buffer was drained from the wells by vacuum, and the worms were subsequently washed once with 600 µL of M9 buffer. After washing, the worms were re-suspended in 50 µL of M9 buffer using low-binding tips, transferred to a new 1.5-mL micro-centrifuge tube, and stored frozen at -80°C. In some cases, the incubation buffer was also collected and frozen at -80°C. For the samples alkylated with iodoacetamide (IAA), worms were resuspended in 50 µL of M9 buffer containing 50 mM iodoacetamide and allowed to incubate for 40 minutes at room temperature before freezing at -80°C. The *P. pacificus* L1 incubations were carried out using the same methods as above, however 100,000 hatchlings were used. The *Rhabditophanes* incubations were carried out similarly, however a mixed population of 50,000 L1/L2 larvae obtained via gravity separation of adults and larvae were used for incubations.

For the wild-type *C. elegans* young adult incubations, a culture of HT115 bacteria harbouring the empty L4440 RNAi feeding vector was grown to an OD_600_ of 0.6 in LB media supplemented with 100 µg/mL ampicillin, and then induced with 1mM IPTG for 1 hour at 20°C until the culture reached an OD_600_ of ∼ 0.8. After induction, the culture was concentrated 5-fold with liquid NGM supplemented with 1 mM IPTG and 100 µg/mL carbenicillin (see ref. 14 for the NGM recipe). 2 mL of this bacterial suspension was added to a 15 mL conical tube. For the treated samples, selectivin-A was added to the suspension to a final concentration of 100 µM and 1% DMSO (v/v). For the untreated control samples, DMSO was added alone to a 1% (v/v) final concentration. 1,250 L1 hatchlings, obtained from an embryo prep of gravid adults^52^, were added to each tube in 0.5 mL of M9 buffer. The worms were incubated on a nutating shaker for 2 days at 25°C until they reached the young adult stage, at which point selectivin-A was added to a final concentration of 100 µM and 1% DMSO (v/v). DMSO alone was added to the untreated control samples. The worms were incubated for 6 hours at 25°C. After incubation, the worms were pelleted by centrifugation, and washed three times with M9 buffer to remove the bacteria. After the final wash, the buffer was aspirated down to 0.5 mL and using low-binding tips the worms were transferred to the wells of a Pall AcroPrep 96-well filter plate (0.45-µm wwPTFE membrane, 1-ml well volume). The buffer was drained from the wells by vacuum. The worms were re-suspended in 50 µL of M9 buffer using low-binding tips, transferred to a new 1.5-mL micro-centrifuge tube, and stored frozen at -80°C. The EMB-8 knockdown incubations were performed identically to the wild-type young adult dose-response experiments, except the MJ69 strain was used in place of the wild-type strain, and worms were fed HT115(DE3) *E. coli* harbouring an RNAi feeding vector expressing dsRNA targeting the *emb-8* gene.

For the *S. cerevisiae* incubations, an overnight culture of the yeast strain RY0568 was diluted to an OD_600_ of 0.1 in 5 mL of YPD and grown at 30°C for 3 hours until the OD_600_ reached 0.3 (see ref. 59 for YPD recipe). This culture was diluted to an OD_600_ of 0.2 in 2.5 mL of YPD and selectivin-A was added at a concentration of 100 µM (1% DMSO (v/v)). DMSO was added alone to the untreated control samples. The culture was incubated for 6h at 30°C until the OD_600_ reached 3.2. The cells were transferred to a 15 mL conical tube and pelleted by centrifugation at 3,000 rpm for 5 minutes. 2 mL of the media was collected and frozen at -80°C, leaving the cells in 0.5 mL of media. The cells were washed twice with distilled and de-ionized water. After the second wash and spin all of the water was removed and the cells were re-suspended in 0.5 mL of M9 buffer. The cells were pelleted again, and all of the M9 buffer was removed by aspiration. The cell pellet was frozen at -80°C.

For the zebrafish incubations, fifty 1-dpf AB wild-type zebrafish were transferred into 0.5 mL of E3 media in 24 well culture plates and allowed to grow to 5-dpf. At 5-dpf, they were transferred into fresh 0.5 mL E3 media in 1.5-mL microcentrifuge tubes and either 100 µM selectivin-A or DMSO alone was added (1% DMSO v/v), as well as 0.05% gelatin to break the surface tension of the water. The fish were incubated for 6 hours at 20°C on a nutating shaker. After the 6-hour incubation, the fish were transferred to the wells of a Pall AcroPrep 96-well filter plate (0.45-µm wwPTFE membrane, 1-mL well volume) using low-binding tips with the ends cut to allow the fish to pass through the opening. The buffer was drained from the wells by vacuum, and the fish were subsequently washed once with 600 µL of M9 buffer. After washing, the fish were re-suspended in 100 µL of M9 buffer using low-binding tips with the ends cut to allow the fish to pass through the opening, and transferred to a new 1.5-mL micro-centrifuge tube. The fish were quickly pelleted by centrifugation and 50 µL of M9 buffer was removed. The fish were then frozen at -80°C. The incubation buffer was also collected and frozen at -80°C.

For the fly larvae incubations, three 3^rd^ instar w1118 fly larvae were transferred into 0.5 mL of PBS buffer in 1.5-mL microcentrifuge tubes. Either 100 µM selectivin-A or DMSO alone was added (1% DMSO v/v). The fly larvae were incubated for 6 hours at 20°C on a nutating shaker. After the 6-hour incubation, the larvae were transferred to the wells of a Pall AcroPrep 96-well filter plate (0.45-µm wwPTFE membrane, 1-mL well volume) using low-binding tips with the ends cut to allow the larvae to pass through the opening. The buffer was drained from the wells by vacuum, and the larvae were subsequently washed once with 600 µL of M9 buffer. After washing, the larvae were transferred into 100 µL of M9 buffer using a fine hair brush. The volume of M9 was adjusted to 50 µL, and the larvae were frozen at -80°C. The incubation buffer was also collected and frozen at -80°C.

### Incubation buffer de-salting and drying

Worm, fish, and fly incubation buffers were de-salted using a 96-well HyperSep C8 solid phase extraction (SPE) plate with 1 mL well volume and 100 mg bed weight (Thermo Scientific). The SPE plates were coupled with a vacuum manifold and vacuum pump, and a flow rate of ∼1 mL per minute was used. The columns were activated with 3 mL of 100% acetonitrile (ACN) and then washed with 3 mL of distilled de-ionized water. Incubation buffers were added to each well and passed through the columns by vacuum. The columns were washed once with 1 mL of water. Three sequential elutions were done using 250 µL of 20% ACN, 250 µL of 50% ACN, and 250 µL of 100% ACN. The eluates were dried in an Eppendorf Vacufuge concentrator, and then stored frozen at -80°C for downstream HPLC analysis.

Yeast incubation buffers were de-salted using Sep-Pak light C8 cartridges (Waters). The columns were activated, loaded, washed, and eluted in the same manner described above for the SPE plates, however solvent and sample was passed through the cartridges using a syringe and manual force. A flow rate of ∼ 1 mL per minute was used. The eluates were dried and stored frozen at -80°C for downstream HPLC analysis.

### Sample lysis for HPLC analysis

#### Worm and fish lysis

Frozen worm and fish pellets were thawed and 50 µL of 2X lysis buffer (20mM Tris-HCl pH 8.3, 0.2% SDS, 240 µg/mL proteinase K) was added to the tubes. The tubes were incubated for 80 minutes in a 56°C water bath with vigorous vortexing every 15 minutes. The samples were then sonicated for 20 minutes at room temperature using a Branson 1510 bath sonicator. The lysates were stored frozen at -80°C until processing by HPLC.

#### Yeast lysis

Frozen yeast pellets were thawed and re-suspended in 40 µL of M9 buffer containing 1M sorbitol and 300 U/mL zymolase. The samples were incubated at 37°C for 60 minutes with vortexing every 10 minutes, after which 50 µL of 2X yeast lysis buffer (20mM Tris-HCl pH 8.3, 0.2% SDS, 720 µg/mL proteinase K) was added to the tubes. The tubes were incubated for 2 hours at 56°C with vigorous vortexing every 10 minutes. The samples were then sonicated for 20 minutes at room temperature using a Branson 1510 bath sonicator. The lysates were stored frozen at -80°C until processing by HPLC.

#### Fly larvae lysis

Frozen larvae were thawed and 50 µL of fly lysis buffer (20mM Tris-HCl pH 8.3, 0.2% SDS, 720 µg/mL proteinase K) was added to the tubes. The larvae were incubated for 30 minutes at 56°C, and were then crushed with a pestle to promote lysis (the exoskeleton remained intact). The samples were incubated at 56°C for an additional 1.5 hours with vigorous vortexing every ten minutes. The samples were then sonicated for 20 minutes at room temperature using a Branson 1510 bath sonicator. The lysates were stored frozen at -80°C until processing by HPLC.

#### Dried incubation buffer preparation

Dried incubation buffers were “lysed” in the same way as the worm pellets, however 50 µL of M9 buffer was added to the dried buffer before proceeding through the lysis steps.

### HPLC method

The frozen lysates were thawed and 50 µL of acidified acetonitrile (ACN) solution (50% ACN, 0.2% acetic acid) was added to the thawed lysates. For the fly samples, three separate lysates were pooled for each replicate to minimize variability associated with having only three larvae per sample. The samples were mixed by vortexing for approximately 10 seconds, and then centrifuged at 17,949*g* for 1 minute. After centrifugation, 50 µL of lysate was injected onto a 4.6 X 150 mm Zorbax SB-C8 column (5 micron particle size) and eluted over 8.65 minutes with the solvent and flow rate gradients shown in Extended Data Table 1. UV-Vis absorbance was measured every 2 nm between 190 and 602 nm. HPLC was performed using an HP 1050 system equipped with an autosampler, vacuum degasser, and variable wavelength diode-array detector. The column was maintained at ∼22°C.

### HPLC quantification

The dry weight of biomaterial in each of the yeast, worm, and fish samples was determined by calculating the area under the curve (AUC) at 300 nm for the major peak of endogenous contents, which elutes between 1 and 1.5 minutes, and then deriving the dry weight in milligrams using standard curves of AUC_300_ vs. dry weight generated using known dry weights for each of the test organisms. The dry weight of fly larvae biomaterial was derived in the same way, however AUC_220_ was used instead of AUC_300_, as it scaled more linearly. To control for differences in biomass, the absorbance intensities of the raw chromatograms, in milliabsorbance units (mAU), were divided by the dry weight of the sample resulting in mAU per mg-of-dry-weight values. Every selectivin-A and 4-(4-chlorophenyl)-1*H*-imidazole-2-thiol/Product 2-treated sample was paired with an untreated DMSO control sample prepared and processed at the same time and in the same way. To correct for background absorbance from endogenous material, the untreated DMSO control mAU per mg-of-dry-weight values were subtracted from the corresponding treated sample mAU per mg-of-dry-weight values to give background-corrected chromatograms. The heat-mapped chromatograms shown in the main figures are these normalized and corrected chromatograms. The heat-mapped chromatograms were generated using custom python scripts. AUC for the analyte peaks was computed using the normalized and corrected chromatograms, and an average across at least three biological replicates was calculated. A peak was considered “not detectable” if an absorbance intensity maximum could not be found at the expected retention time. If a peak was not detectable it was given an AUC value of zero. AUCs for M1, M2, M3 and P2-alk were calculated at 260 nm, AUC for M4 was calculated at 300 nm. The same quantification scheme was used for the incubation buffer samples, with two minor exceptions for the yeast incubation buffers: 1. A YPD blank sample prepared in the same way as the experimental yeast buffers was processed by HPLC and the absorbance intensity values were subtracted from the experimental intensities to account for absorbing material from the YPD media – this was done prior to dry weight normalization and subtraction of the paired DMSO control absorbance intensities; 2. Since only 2 out of 2.5 mL (4/5) of yeast incubation buffer was processed, the dry weight values used to normalize the yeast buffer data were multiplied by 4/5. Average AUC values were calculated across at least three experimental replicates. Microsoft Excel was used for all chromatogram normalizations, background corrections, and AUC determinations.

### Metabolite mass spectrometry

HPLC fractions containing the selectivin-A metabolites M1, M2, M3, M4, and P2-alk, were collected from three separate lysates, combined, and dried using a SpeedVac concentrator. The identical fractions from DMSO control lysates were also collected and dried. The dried fractions were re-suspended in a minimal volume of 1:1 (*v/v*) methanol: 0.1% aqueous formic acid. Electrospray ionization mass spectrometry (ESI-MS) analyses were carried out using a 6538 UHD model quadrupole time-of-flight mass analyzer equipped with an atmospheric pressure ESI source and a 1260 Infinity model HPLC system (Agilent Technologies, Santa Clara, CA). Samples were analyzed via loop injection with mobile phase composed of 1:1 (*v/v*) methanol: 0.1% aqueous formic acid and flowing at a rate of 0.25 mL min^-1^. Mass spectra were recorded in the 2 GHz mode and the high-resolution MS analyses for molecular formula determinations were obtained using external calibration. Tandem MS/MS analyses were obtained via collision-induced dissociation using the targeted MSn function of the acquisition software. MS/MS spectra were recorded sequentially at three different fragmentation voltages (10, 20 and 30 V) and the resulting spectrum was composed of the average of those three collision energies.

### Attempted oxidation of selectivin-A

In an attempt to generate the sulfoxide metabolite of selectivin-A in the lab, we tried three different oxidation methods: 1. Hydrogen peroxide as oxidant: H_2_O_2_ (1 equiv), acetone/H_2_O (5:1) 0°C to RT; 2. Oxone as oxidant: oxone (5 equiv), H_2_O/MeOH, RT; 3. m-CPBA as oxidant: m-CPBA (2.5 equiv), DCM, RT. Using hydrogen peroxide as the oxidant resulted in the recovery of starting material alone (i.e. selectivin-A), and cleavage of the imidazole ring occurred when oxone was used as the oxidizing agent. When m-CPBA was used as the oxidant, a complex mixture was observed and purification only resulted in further decomposition.

### Synthesis of selectivin-E, -F, -H, and -I

For each selectivin analog, the appropriate 2-bromo-acetophenone was synthesized from the corresponding commercially available acetophenone according to literature procedures^60^. The imidazo[2,1-*b*]thiazoles were prepared according to a modified literature procedure^61^. To a 2 dram vial was added the appropriate α-bromoketone (1 mmol, 1 equiv), the appropriate 2-aminothiazole (1.3 mmol, 1.3 equiv), and EtOH (1.6 mL) and the reaction mixture was stirred at reflux until disappearance of the α-bromoketone was evident by TLC. The mixture was concentrated, then purified by column chromatography using the given eluent to provide the imidazo[2,1-*b*]thiazole.

#### 6-(4-chlorophenyl)-2-methylimidazo[2,1-b]thiazole (selectivin-E)

10 mmol (2.3g) scale. Purified using pentanes–EtOAc (15:5 to 10:10 v:v). Pale orange solid (30%). ^1^H-NMR (CDCl_3_, 500 MHz): 7.73 (d, *J* = 8.5 Hz, 2H), 7.59 (s, 1H), 7.34 (d, *J* = 8.5 Hz, 2H), 7.13 (q, *J* = 1.4 Hz, 1H), 2.42 (d, *J* = 1.4 Hz, 3H). ^13^C{^1^H}-NMR (CDCl_3_, 125 MHz): 149.9, 145.5, 132.9, 132.8, 128.9, 127.0, 126.4, 115.2, 107.9, 14.2.

#### 6-(4-fluorophenyl)-3-methylimidazo[2,1-b]thiazole (selectivin-F)

Purified using pentanes–EtOAc (15:5 v:v). Brown solid (38%, MP = 109-114 °C). ^1^H-NMR (CDCl_3_, 500 MHz): 7.83 – 7.78 (m, 2H), 7.57 (s, 1H), 7.12 – 7.05 (m, 2H), 6.42 (q, *J* = 1.3 Hz, 1H), 2.43 (d, *J* = 1.3 Hz, 3H). ^13^C{^1^H}-NMR (CDCl_3_, 125 MHz): 162.4 (d, *J* = 246.2 Hz), 149.9, 146.9, 130.4, 127.9, 127.0 (d, *J* = 8.0 Hz), 115.7 (d, *J* = 21.6 Hz), 107.0, 105.8, 13.5. ^19^F{^1^H}-NMR (CDCl_3_, 375 MHz): -115.0. IR (neat): 3134, 2965, 2926, 2883, 1750, 1475, 1375, 1155, 1092, 1009, 831, 755, 692. Mass: DART+, calc. for C_12_H_10_N_2_FS 233.05432 [M+H]^+^, found 233.05424.

#### 6-(4-bromophenyl)-3-methylimidazo[2,1-b]thiazole (selectivin-H)

Purified using pentanes–EtOAc (16:4 to 15:5 v:v). Orange solid (33%). The spectral data were in accordance with literature^62^. ^1^H-NMR (CDC_l3_, 500 MHz): 7.73 – 7.69 (m, 2H), 7.61 (s, 1H), 7.53 – 7.49 (m, 2H), 6.42 (q, *J* = 1.3 Hz, 1H), 2.42 (d, *J* = 1.3 Hz, 3H). ^13^C{^1^H}-NMR (CDCl_3_, 125 MHz): 150.1, 146.8, 133.4, 131.9, 127.8, 126.8, 121.2, 107.1, 106.3, 13.5.

#### 6-(4-bromophenyl)-2-methylimidazo[2,1-b]thiazole (selectivin-I)

Purified using pentanes–EtOAc (16:4 to 8:12 v:v). White solid (32%, MP = 235-240 °C). ^1^H-NMR (CDCl_3_, 500 MHz): 7.69 – 7.64 (m, 2H), 7.61 (s, 1H), 7.52 – 7.47 (m, 2H), 7.13 (q, *J* = 1.4 Hz, 1H), 2.42 (d, *J* = 1.5 Hz, 3H). ^13^C{^1^H}-NMR (CDCl_3_, 125 MHz): 150.0, 145.5, 133.2, 131.9, 127.0, 126.7, 121.0, 115.2, 108.0, 14.2. IR (neat): 3134, 2965, 2926, 2883, 1750, 1475, 1375, 1155, 1092, 1009, 831, 755, 692. Mass: DART+, calc. for C_12_H_10_N_2_SBr 292.97426 [M+H]^+^, found 292.97416.

Work-up and isolation of compounds was performed using standard benchtop techniques. All commercial reagents were purchased from chemical suppliers (Sigma-Aldrich, Combi-Blocks, Alfa Aesar, or Strem Chemicals) and used without further purification. Dry solvents were obtained using standard procedures (THF was distilled over sodium/benzophenone, dichloromethane was distilled over calcium hydride). Reactions were monitored using thin-layer chromatography (TLC) on EMD Silica Gel 60 F254 plates. Visualization was performed under UV light (254nm) or using potassium permanganate (KMnO_4_) or I_2_ stain. Flash column chromatography was performed on Siliaflash P60 40-63 µm silica gel purchased from Silicycle. NMR characterization data was obtained at 293K on a Varian Mercury 300 MHz, Varian Mercury 400 MHz, Bruker Advance III 400 MHz, Agilent DD2 500 MHz equipped with a 5mm Xses cold probe or Agilent DD2 600 MHz. ^1^H spectra were referenced to the residual solvent signal (CDCl_3_ = 7.26 ppm, DMSO-*d_6_* = 2.50 ppm). ^13^C{^1^H} spectra were referenced to the residual solvent signal (CDCl_3_ = 77.16 ppm, DMSO-*d_6_* = 39.52 ppm). Data for ^1^H NMR are reported as follows: chemical shift (δ ppm), multiplicity (s = singlet, d = doublet, t = triplet, q = quartet, m = multiplet), coupling constant (Hz), integration. NMR spectra were recorded at the University of Toronto Department of Chemistry NMR facility. Infrared spectra were recorded on a Perkin-Elmer Spectrum 100 instrument equipped with a single-bounce diamond/ZnSe ATR accessory in the solid state and are reported in wavenumber (cm^-1^) units. Melting point ranges were done on a Fisher-Johns Melting Point Apparatus and are reported uncorrected. High resolution mass spectra (HRMS) were recorded at the Advanced Instrumentation for Molecular Structure (AIMS) in the Department of Chemistry at the University of Toronto.

### Meloidogyne incognita tomato plant root infection experiments

An *M. incognita* population originally collected from grape (*Vitis vinifera*) in Parlier, California, was used in all experiments, and they were maintained on tomato plants (*Solanum lycopersicum* ‘Rutgers’) as previously described^56^. Infective *M. incognita* second-stage juveniles (J2s) were collected as described in^56^. For the infection assays, 90 grams of soil (1:1 sand:loam mix) was added to each compartment of several 6-compartment plastic garden packs. The soil was drenched with 18 mL of deionized water containing dissolved chemical or DMSO alone. Depending on the batch, either 1,000 or 2,500 infective J2s were then added to the soil in 2 mL of water, for a total water volume of 20 mL. All of the tests done in the same batch and on the same day had the same number of J2s added to the soil so that the DMSO control and treatment samples had the same number of J2s added. The final aqueous DMSO concentration varied from 0.05% to 0.4% (v/v) depending on the stock concentration of the chemical and the final concentration of the chemical tested. The highest DMSO concentration of 0.4% (v/v) was used for all DMSO controls. Importantly, we observed that 0.4% DMSO did not inhibit root infection compared with water control. There were four DMSO controls in each batch. The J2s were incubated in the soil and chemical for 24 hours, after which two-to three-week old tomato seedlings were transplanted into the soil (one plant per compartment). Inoculated plants were grown for 8 weeks in a greenhouse as described^56^, under long-day conditions (16-h photoperiod) with 26/18°C day/ night temperatures. After 8 weeks, the plants were destructively harvested. The tops were removed and discarded, and roots were gently washed with water to remove adhering soil. Eggs were extracted by placing rinsed roots in 0.6% sodium hypochlorite and agitating at 300 rpm for 3 min. Roots were then rinsed over nested 250- and 25.4-µm sieves, with eggs collected from the latter and suspended in water. Roots were dried in a 65°C oven for at least 72 hours, after which dry roots were weighed. The number of eggs from each plant root was counted on a dissection microscope using a haemocytometer, and the number of eggs per milligram of root was calculated by dividing the total egg number by the mass of the dried root material. The eggs per milligram of root value for each treatment replicate was normalized to the average of the four DMSO controls for a given batch. To calculate percent effectiveness, the normalized values were subtracted from 1, and then multiplied by 100. An average percent effectiveness value was then calculated across at least four replicates (no eggs = 100% effectiveness; same number of eggs relative to control = 0% effectiveness; more eggs relative to control = less than 0% effectiveness). Parts-per-million soil concentrations were calculated as µg active ingredient per gram of soil. Killograms-per-hectare values were calculated assuming a soil depth of 15 cm and a soil density of 1200 kg/m^3^.

### Forward genetic screens for selectivin-resistant mutants

Forward genetic screens were carried out as previously described^14^. Briefly, wild-type parental (P0) worms were mutagenized in 50mM ethyl methanesulfonate (EMS) for 4 hours. Synchronized L1s from either the F1 (progeny) or F2 (grand-progeny) generations were dispensed onto 10cm MYOB agar plates (see ref. 52 for how to prepare MYOB/agar media) containing a 100% penetrant lethal dose of the nematicide. Worms were plated at a density of 20,000 L1s per plate. The plates were scanned by eye for viable worms.

### Statistical information

HPLC analyses of unmodified selectivin parent and metabolites were performed at least three times for each strain or organism, and area under the curve (AUC) for each analyte was calculated and averaged across the replicates. Data are presented as means ± SEM. P-values were calculated using unpaired one-tailed Student’s t-tests assuming equal SD. For the selectivin analog series, soil-based root infection assays were performed at least 4 times. Ordinary one-way ANOVA was used to estimate differences between the means for the 15 compounds assayed (F = 6.441, DFn = 14, DFd = 69, P < 0.0001). Uncorrected Fisher’s LSD was then used to calculate a P-value for each pairwise comparison of means (P > 0.05 suggests that the means are not significantly different). No statistical methods were used to predetermine sample size. The investigators were not blinded to allocation during experiments and outcome assessment. GraphPad prism was used for all statistical analyses.

### Data Availability

All data that support the conclusions of this manuscript are available upon request.

## ACKNOWLEDGEMENTS

Strains were provided by the *Caenorhabditis* Genetics Center (University of Minnesota) and Marie-Anne Félix (IBENS, Paris, France). Mass spectrometry analysis was performed by the AIMS Mass Spectrometry Laboratory in the Department of Chemistry at the University of Toronto. Many thanks to Dr. Doug Colwell and Dawn Gray at the Lethbridge and Agri-Food Canada Research Centre for supplying faeces from infected calves for *Cooperia oncophora* egg collection. We thank Tim Hughes for the use of their SpeedVac, Brian Ciruna for the gift of NACET, Grant Brown for the culture of *S. cerevisiae*, Brent Derry for helpful comments on the manuscript, and Nicole Robbins from Leah Cowen’s lab for comments on the work.

## AUTHOR CONTRIBUTIONS

Unless otherwise noted all experiments were carried out in the lab of P.J.R. All free-living nematode dose-response experiments were carried out by A.R.B. A.R.B performed the HPLC-based metabolism experiments, and analyzed all related mass spectrometry data. A.R.B. performed the NACET experiments. B.M.P., E.M.R., and A.R.B. performed the *Cooperia* dose-response experiments in J.S.G.’s lab. *S. cerevisiae* experiments were done by A.R.B. J.S. performed the HEK cell dose-response assays in I.S.’s lab. J.T. in the lab of H.M.K., as well as J.R.V. in the lab of J.J.D., performed the zebrafish dose-response assays. S.M. and S.W.C. performed the HepaRG dose-response experiments in S.A.M.’s lab. E.P. performed the HepG2 and *C. albicans* assays in L.E.C.’s lab. In the lab of S.R.C., A.S.V. performed the *Arabidopsis* greening assays. C.A.M.F. and Q.X. carried out the mouse experiments. R.J.R. carried out the selectivin syntheses in M.L.’s lab. M.H.M. carried out the *in vitro M. incognita* assays in S.L.F.M.’s lab. M.K. carried out the *in vitro M. chitwoodi* assays as well as the *M. incognita* soil-based reproduction assays in tomato plants in the lab of I.Z. *D. melanogaster* experiments were carried out by C.H. and A.R.B, in the labs of P.J.R. and H.M.K. The genetic screens for resistant mutants were performed by A.R.B. The project was conceived by A.R.B. and P.J.R. The manuscript was written by A.R.B. and P.J.R.

## ADDITIONAL INFORMATION

**Supplementary Information** is available for this paper.

Correspondence and requests for materials should be addressed to Peter J. Roy or Andrew R. Burns.

## EXTENDED DATA FIGURE AND TABLE LEGENDS

**Extended Data Fig. 1.**
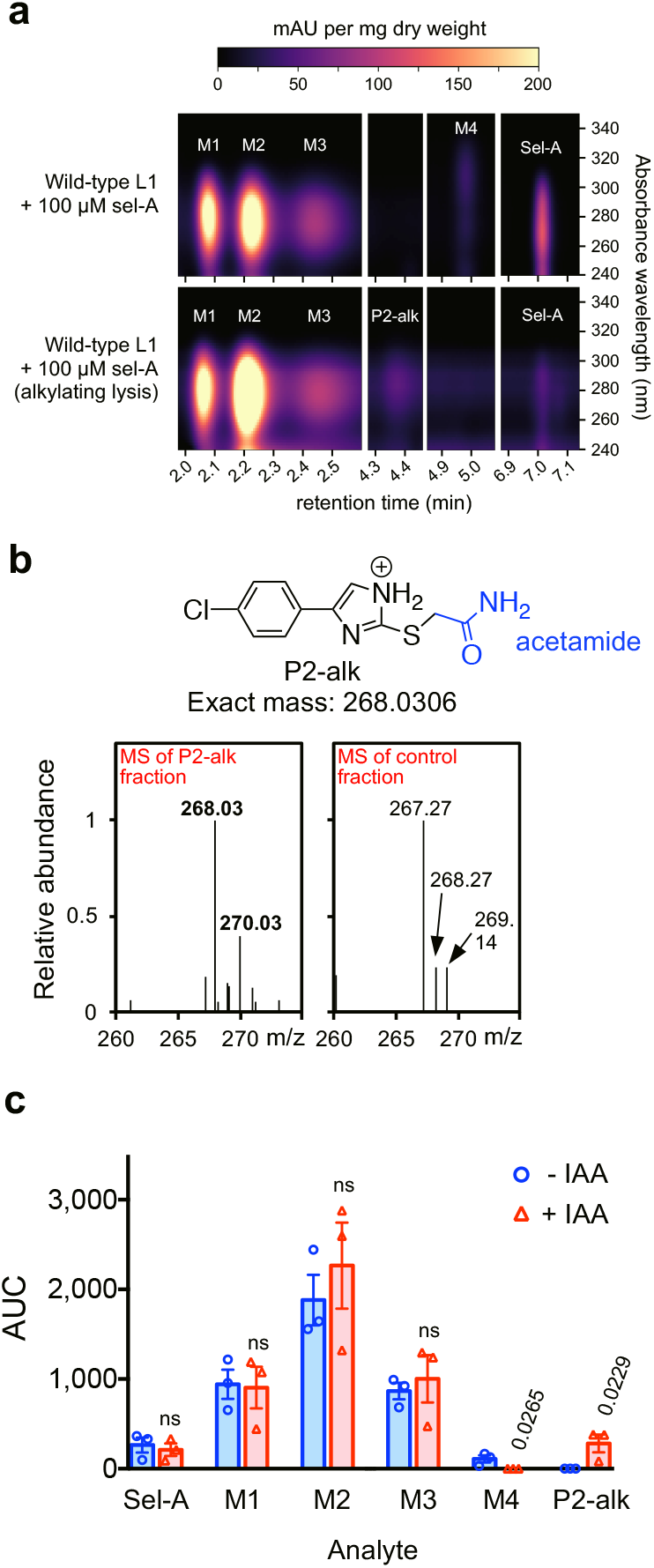
The selectivin-A imidazole-thiol metabolite and the low-molecular-weight thiol conjugates of the sulfoxide metabolite are produced *in vivo*. **a,** Biomass-normalized and background-corrected HPLC chromatograms for lysates of selectivin-A-treated L1 worms lysed normally or in the presence of the alkylating agent iodoacetamide (IAA). The peaks corresponding to unmodified selectivin-A and metabolites M1, M2, M3, M4, and alkylated Product 2 (P2-alk) are indicated. **b,** Mass spectrometry data for the P2-alk HPLC fraction and the corresponding untreated control. **c,** Quantification of selectivin-A, M1, M2, M3, M4, and P2-alk in the lysates of selectivin-A-treated L1 worms either lysed normally or in the presence of IAA. AUC is area under the curve for the given analyte peak. AUCs for the M1, M2, M3, and P2-alk metabolites were calculated at 260 nm. AUC for the M4 metabolite was calculated at 300 nm. Error bars are SEM. For each analyte, unpaired one-tailed Student’s t-tests were performed comparing the means of the plus or minus IAA conditions. P-values > 0.05 are not significant (ns). P-values < 0.05 are shown.

**Extended Data Fig. 2.**
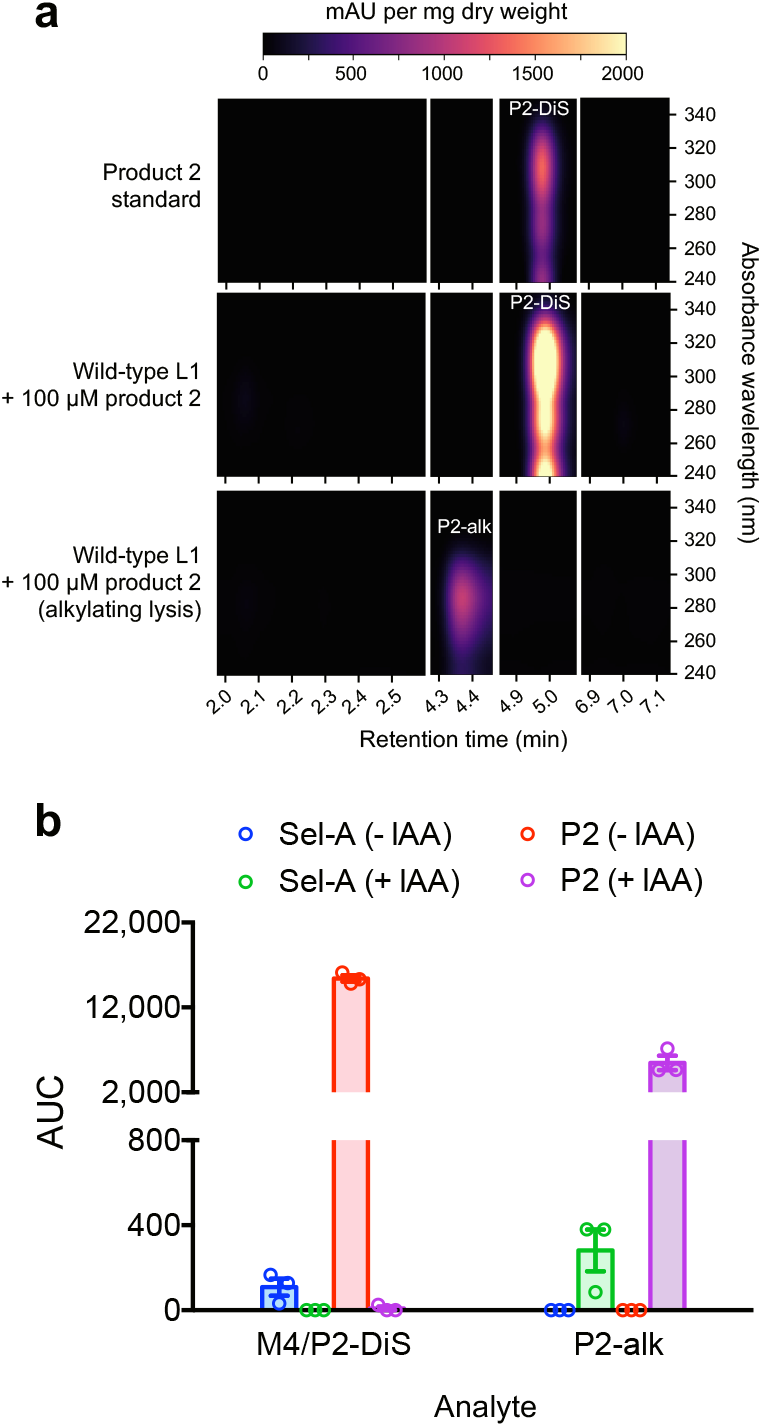
Selectivin metabolic product 2 is bioavailable to worms when applied exogenously. **a,** Biomass-normalized and background-corrected HPLC chromatograms for lysates of L1 worms incubated in product 2 (P2), lysed normally or in the presence of iodoacetamide (IAA). The top chromatogram is the product 2 (P2) standard injected directly onto the column (this chromatogram is not biomass-normalized or background-corrected). The P2 standard oxidizes to form the disulfide (P2-DiS) prior to HPLC analysis. The disulfide (P2-DiS) and alkylated (P2-alk) peaks are indicated. **b,** Quantification of M4/P2-DiS and P2-alk in the lysates of 100 µM selectivin-A-treated or 100 µM P2-treated L1 worms, lysed normally or in the presence of IAA. AUC is area under the curve for the given analyte peak and was calculate at 260 nm. Error bars are SEM. The y-axis is broken between 800 and 2000 to accommodate the large amounts of P2 in the lysates of worms.

**Extended Data Fig. 3.**
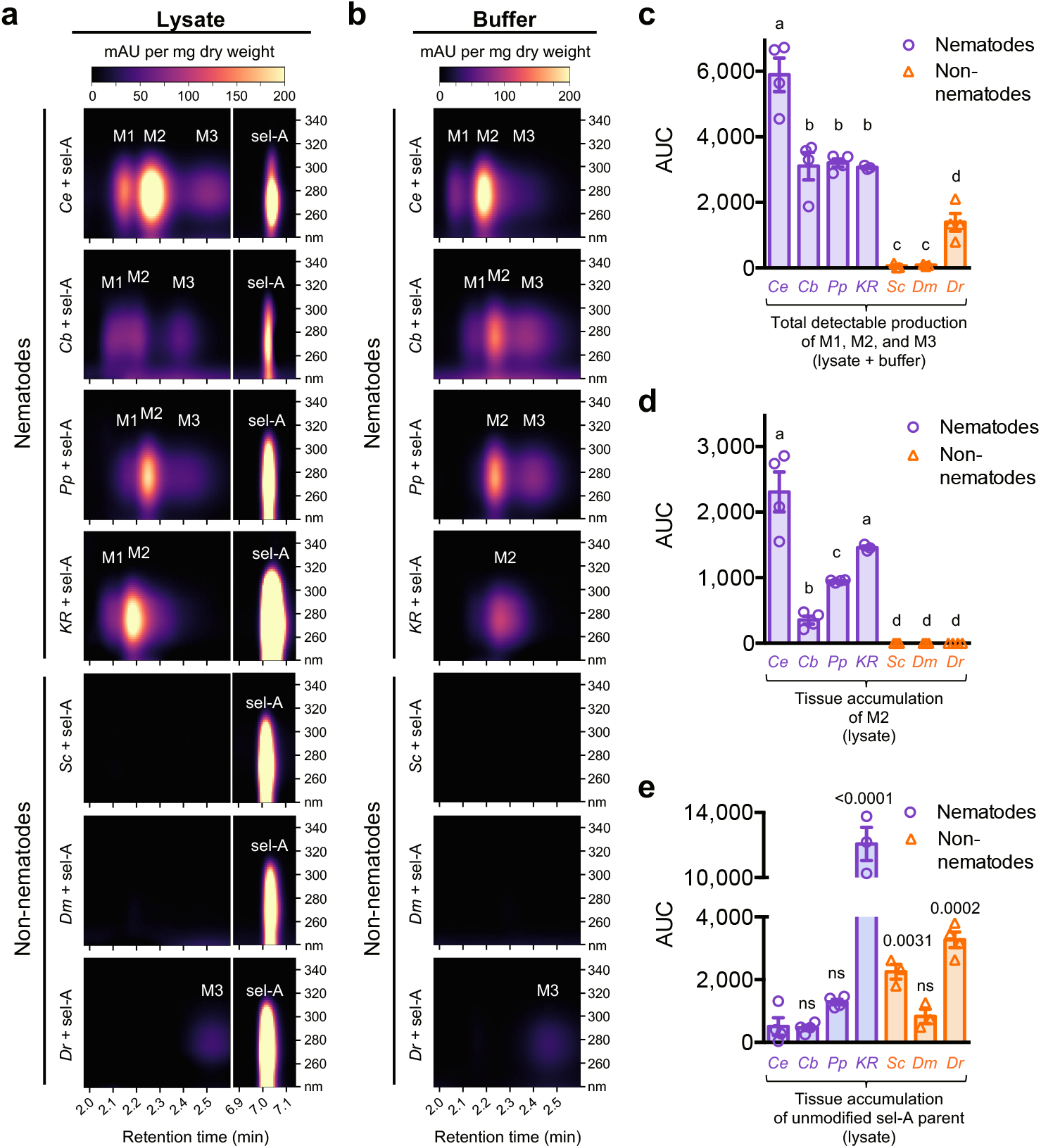
Selectivin bioactivation is nematode-specific. **a and b,** Biomass-normalized and background-corrected HPLC chromatograms of lysates (**a**) and incubation buffers (**b**) from *C. elegans* (*Ce*), *C. briggsae* (*Cb*), *P. pacificus* (*Pp*), *Rhabditophanes sp.* KR3021 (*KR*), *D. melanogaster* (*Dm*), and *D. rerio* (*Dr*) larvae, as well as *S. cerevisiae* (*Sc*) cells, incubated in 100 µM selectivin-A. Selectivin-A and metabolite peaks are indicated. **c,** Total detectable production of M1+M2+M3 in lysate and buffer for each of the seven species tested. **d,** M2 tissue accumulation in each species tested. **e,** Unmodified selectivin-A tissue accumulation in each of the test species. AUC is area under the curve at 260 nm. Error bars are SEM. For **c** and **d**, unpaired one-tailed Student’s t-tests were performed for all pairwise comparisons of means. The means sharing a letter are statistically indistinguishable (p > 0.01). For **e**, unpaired one-tailed Student’s t-tests were performed relative to the *C. elegans* mean. P-values > 0.01 are not significant (ns). P-values < 0.01 are shown.

**Extended Data Fig. 4.**
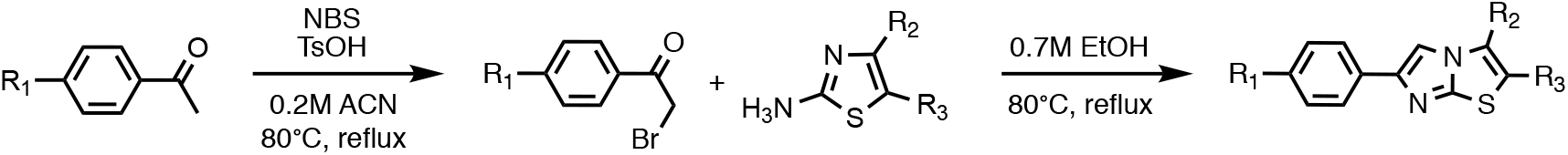
Selectivin synthesis follows a simple synthetic route. Synthesis of the selectivins is a simple two-step procedure that starts with the α-bromination of the commercially available acetophenones followed by condensation with the appropriate heterocycle. The starting materials are all inexpensive and readily purchasable, and the synthetic steps make use of common reagents and solvents such as N-bromosuccinimide (NBS), p-toluenesulfonic acid (TsOH), acetonitrile (ACN), and ethanol (EtOH). The synthesis is performed using moderate reaction temperatures, and reasonably high yields can be achieved without isolation or purification of the reaction intermediates.

**Extended Data Fig. 5.**
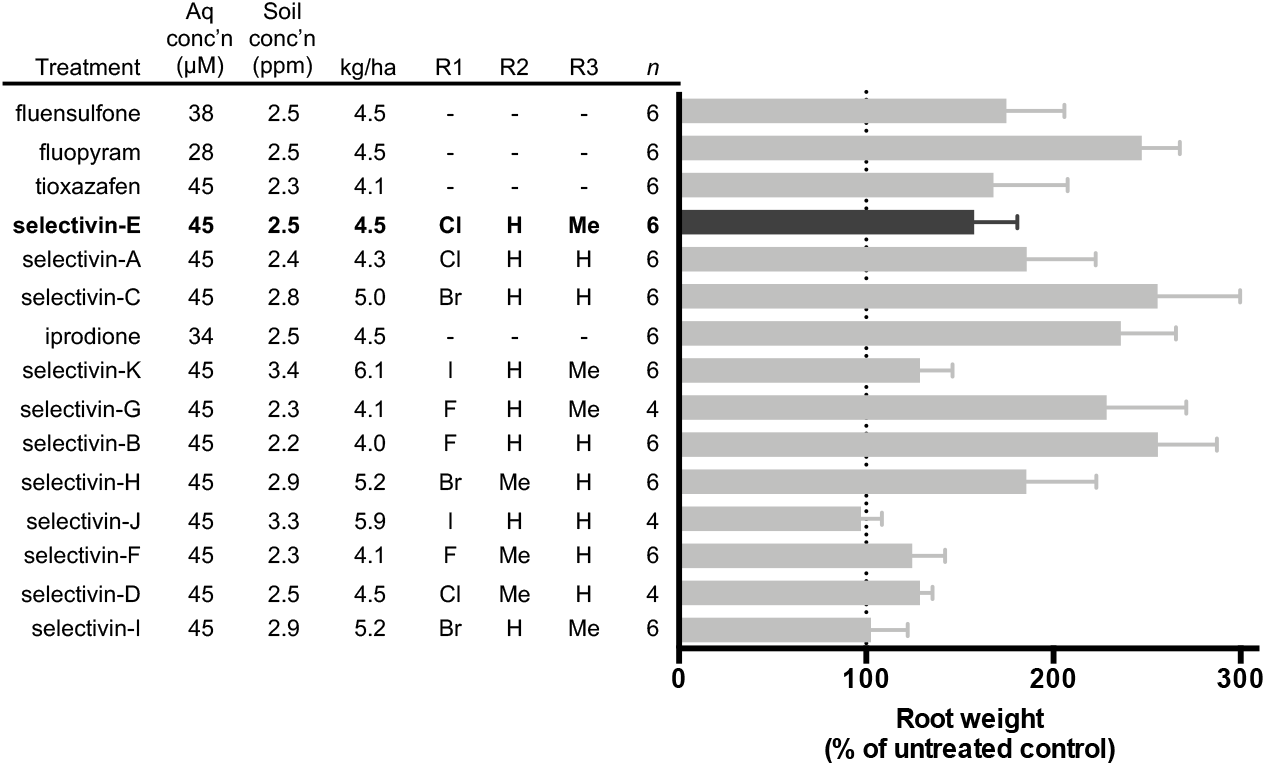
Selectivin treatment generally increases the root weights of tomato plants challenged with nematode infection. The effects of 11 selectivin analogs and 4 commercial nematicides on the root weights of tomato plants challenged with *M. incognita* infection is shown. Root weight values are percent of untreated control. For each analog, aqueous molar concentration, parts-per-million soil concentration, and kilograms-per-hectare values are shown. The R-groups for each selectivin analog are indicated (see Fig. 1a for the selectivin core scaffold and R-group positions).

**Extended Data Table 1.** Anthelmintic/nematicide-resistant mutants are sensitive to the selectivins. Selectivin-A and selectivin-C LC50 data for wild-type worms and nine distinct nematicide/anthelmintic-resistant mutant strains. LC50 is the concentration at which 50% of the nematodes are dead. OP, organophosphate; LEV, levamisole; APA, aminophenylamidine; THP, tetrahydropyrimidine; AAD, aminoacetonitrile derivative; BZ, benzimidazole.

| Genotype | Resistance | LC50 (μM) | |
| --- | --- | --- | --- |
|  |  | selectivin-A | selectivin-C |
| wild-type | None | 6.9 | 3.6 |
| *cha-1(p1152)* | Carbamates/OPs | 3.7 | 2.3 |
| *avr-14(ad1305);*<br>*avr-15(vu227); glc-1(pk54)* | Ivermectin/<br>Abamectin | 4.6 | 3.0 |
| *mev-1(tr393)* | Fluopyram | 4.9 | 1.6 |
| *unc-63(ok1075)* | LEV, APAs, THPs | 6.4 | 3.1 |
| *acr-23(ok2804)* | AADs | 6.9 | 3.3 |
| *daf-16(mu86)* | Apigenin | 5.7 | 3.1 |
| *ben-1(e1880)* | BZs | 6.4 | 3.1 |
| *slo-1(js379)* | Emodepside | 4.4 | 3.1 |

**Extended Data Table 2.** Genetic resistance to selectivins is difficult to achieve. The number of genomes screened and the number of resistant mutants obtained from genetic screens for mutants that resist selectivin-A, selectivin-C, and four additional commercial nematicides/anthelmintics. Wact-11 is a structural analog of the commercial nematicide fluopyram. Wact-11 and fluopyram share the same nematicidal mode-of-action, i.e. inhibition of mitochondrial complex II, and wact-11 resistant mutants are also resistant to fluopyram^14^.

| Nematicide/<br>anthelmintic | # of F1<br>genomes<br>screened | # of F2<br>genomes<br>screened | # of resistant<br>mutants obtained | Reference |
| --- | --- | --- | --- | --- |
| selectivin-A | 10 million | 100,000 | 0 | This study |
| selectivin-C | 150,000 | 50,000 | 0 | This study |
| levamisole | 0 | 10,000 | 31 | Ref. 24 |
| albendazole | 0 | 10,000 | 22 | Ref. 24 |
| ivermectin<br>(abamectin analog) | 0 | 10,000 | 8 | Ref. 24 |
| wact-11<br>(fluopyram analog) | 1.3 million | 180,000 | 15 | Ref. 14 |

**Extended Data Table 3.**
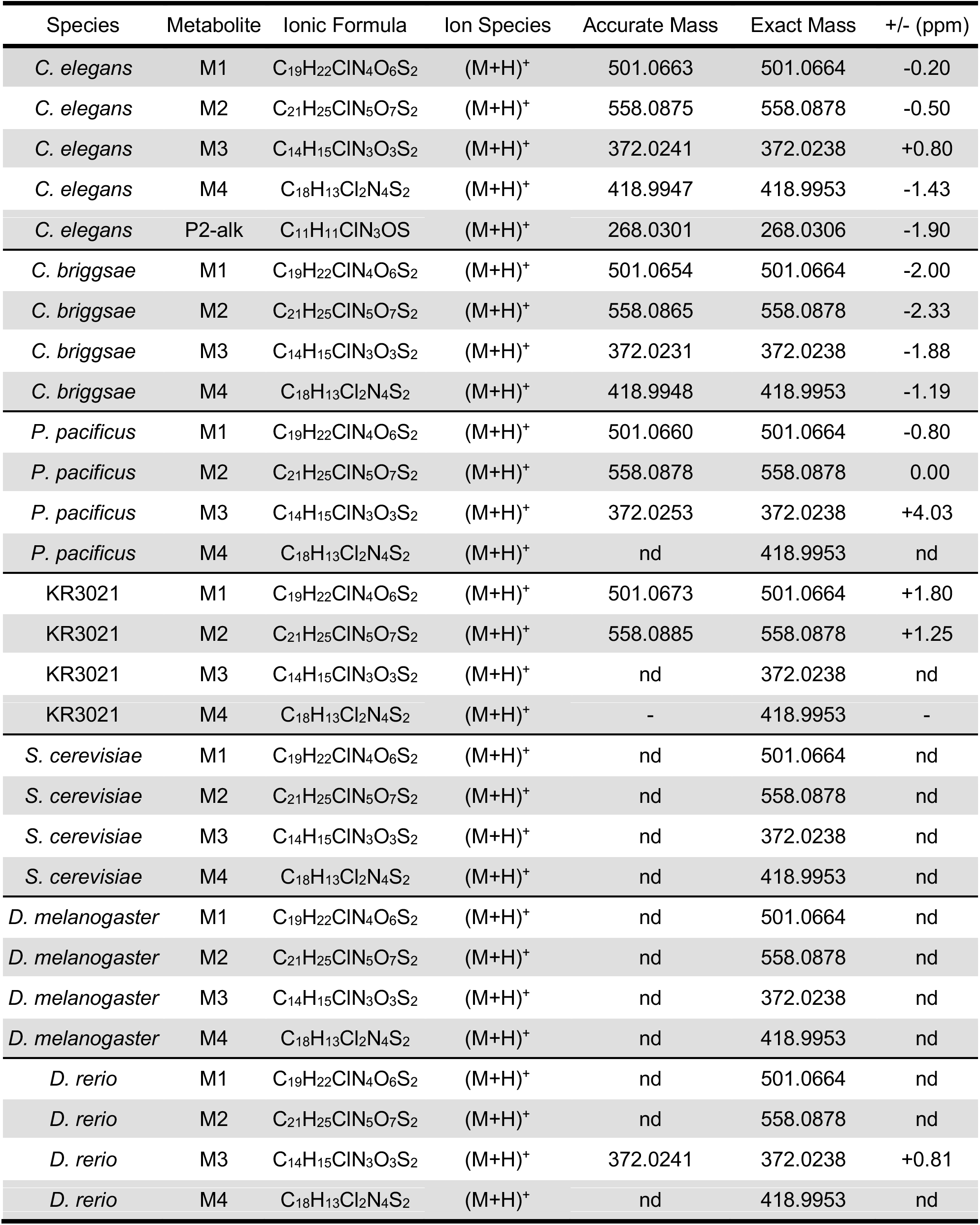
Accurate mass and exact mass values for the selectivin-A metabolites. Accurate mass, exact mass, and the parts-per-million (ppm) difference between the two masses for the proposed selectivin-A metabolites for each species tested. A difference of less than 5 ppm between the accurate mass determined by MS and the exact mass of the proposed metabolites indicates that the molecular formula is correct. KR3021 is *Rhabditophanes sp.* KR3021. The KR3021 M4 fraction was not processed for technical reasons. The abundance of some masses was too low in the mass spectrum to perform accurate mass determinations, and so they were not determined (nd).

**Extended Data Table 4.** Summary of MS data for selectivin-A metabolites from lysates. Raw MS counts for the m/z values corresponding to the selectivin-A (sel-A) metabolites M1, M2, M3, and M4 in HPLC fractions taken from lysates of four nematode species and three non-nematode species incubated in selectivin-A. Raw counts were taken from centroid plots of the raw mass spectra. Counts below 1 x 10^3^ were considered not detectable (nd). Ce, *C. elegans*; Cb, *C. briggsae*; Pp, *P. pacificus*; KR, *Rhabditophanes sp*. KR3021; Sc, *S. cerevisiae*; Dm, *D. melanogaster*; Dr, *D. rerio*. For technical reasons, the KR M4 metabolite fraction was not processed by MS.

| Incubation Condition | Metabolite Fraction | m/z | Raw Counts |  |  |  |  |  |  |
| --- | --- | --- | --- | --- | --- | --- | --- | --- | --- |
|  |  |  | Ce | Cb | Pp | KR | Sc | Dm | Dr |
| + sel-A | M1 | 501.07 | 72,051 | 102,049 | 31,398 | 19,552 | nd | nd | 1,978 |
| + sel-A | M2 | 558.09 | 484,191 | 249,692 | 325,518 | 163,260 | nd | nd | 4,862 |
| + sel-A | M3 | 372.02 | 81,766 | 24,753 | 19,842 | 3,800 | nd | 1,211 | 8,495 |
| + sel-A | M4 | 418.99 | 39,766 | 53,196 | nd | - | nd | nd | 5,102 |
| - sel-A | M1 | 501.07 | nd | nd | nd | nd | nd | nd | nd |
| - sel-A | M2 | 558.09 | nd | nd | nd | nd | nd | nd | nd |
| - sel-A | M3 | 372.02 | nd | nd | nd | nd | nd | nd | nd |
| - sel-A | M4 | 418.99 | nd | nd | nd | - | nd | nd | nd |

**Extended Data Table 5.** HPLC solvent and flow rate gradients. Solvent and flow rate gradients for the HPLC method used in this study. Solvent A is 4.9:95:0.1 (ACN:H_2_O:acetic acid). Solvent B is 95:4.9:0.1 (ACN:H_2_O:acetic acid).

| **Time** | **Flow Rate (ml/min)** | **% Solvent A** | **% Solvent B** |
| --- | --- | --- | --- |
| 0.00 | 1.3 | 85.0 | 15.0 |
| 4.49 | 1.3 | 43.5 | 56.5 |
| 4.50 | 1.0 | 43.5 | 56.5 |
| 5.54 | 1.0 | 42.0 | 58.0 |
| 6.54 | 2.0 | 30.0 | 70.0 |
| 8.04 | 3.0 | 0.0 | 100 |
| 8.34 | 3.0 | 0.0 | 100 |
| 8.35 | 2.0 | 85.0 | 15.0 |

